# FLIP(L) determines p53 induced life or death

**DOI:** 10.1101/858688

**Authors:** Andrea Lees, Alexander J. McIntyre, Fiammetta Falcone, Nyree T. Crawford, Christopher McCann, Gerard P. Quinn, Jamie Z. Roberts, Tamas Sessler, Peter F. Gallagher, Gemma M.A. Gregg, Katherine McAllister, Kirsty M. McLaughlin, Wendy L. Allen, Caitriona Holohan, Laurence J. Egan, Aideen E. Ryan, Melissa Labonte-Wilson, Phillip D. Dunne, Mark Wappett, Vicky M. Coyle, Patrick G. Johnston, Emma M. Kerr, Daniel B. Longley, Simon S. McDade

**Affiliations:** Centre for Cancer Research and Cell Biology, Queen’s University Belfast, Belfast, Northern Ireland, BT9 7BL, UK; Regenerative Medicine Institute (REMEDI), College of Medicine, Nursing and Health Sciences, NUI Galway, Galway, Ireland; Discipline of Pharmacology & Therapeutics, Lambe Institute for Translational Research, School of Medicine, College of Medicine, Nursing and Health Sciences, NUI Galway, Galway, Ireland

## Abstract

How p53 differentially activates cell cycle arrest versus cell death remains poorly understood. Here, we demonstrate that upregulation of canonical pro-apoptotic p53 target genes in colon cancer cells imposes a critical dependence on the long splice form of the caspase-8 regulator FLIP (FLIP(L)), which we identify as a direct p53 transcriptional target. Inhibiting FLIP(L) expression with siRNA or Class-I HDAC inhibitors promotes apoptosis in response to p53 activation by the MDM2 inhibitor Nutlin-3A, which otherwise predominantly induces cell-cycle arrest. When FLIP(L) upregulation is inhibited, apoptosis is induced in response to p53 activation via a novel ligand-independent TRAIL-R2/caspase-8 complex, which, by activating BID, induces mitochondrial-mediated apoptosis. Notably, FLIP(L) depletion inhibits p53-induced expression of the cell cycle regulator p21 and enhances p53-mediated upregulation of PUMA, with the latter activating mitochondrial-mediated apoptosis in FLIP(L)-depleted, Nutlin-3A-treated cells lacking TRAIL-R2/caspase-8. Thus, we report two previously undescribed, novel FLIP(L)-dependent mechanisms that determine cell fate following p53 activation.

## Introduction

The mechanisms through which p53 activation differentially regulates cell survival (*e.g.* by initiating cell cycle arrest and DNA damage repair) and cell death (by promoting apoptosis) remain enigmatic^1^. Canonically, phosphorylation of the p53 N-terminus blocks its interaction with the E3 ubiquitin ligase Mouse Double Minute 2 (MDM2) (Momand *et al*, 1992; Shieh *et al*, 1997) reducing ubiquitination of the C-terminus and stabilizing p53. Stabilized p53 directly activates target genes to induce cell cycle arrest mediated by the cyclin-dependent kinase inhibitor p21/*CDKN1A* and other effectors (el-Deiry *et al*, 1993; Harper *et al*, 1993). In the event of sustained stress and/or irreparable DNA damage, sequential post-translational modification of p53 is thought to direct p53 activity towards pro-apoptotic target genes, including those encoding Fas/CD95 (*FAS*) (Müller *et al*, 1998), TRAIL Receptor 2 (TRAIL-R2/*TNFRSF10B*) also known as death receptor-5 (DR5) (Takimoto & el-Deiry, 2000), Bax/*BAX* and the BH3 proteins Puma/*BBC3* and Noxa/*PMAIP1* (Nakano & Vousden, 2001; Villunger, 2003). While we (McDade *et al*, 2014) and others have contributed to progress in identifying direct and indirect p53 target genes using ChIP-seq (Fischer *et al*, 2016), the mechanisms underlying the switch from cell cycle arrest to cell death induction remains poorly understood.

Despite p53 being the most frequently mutated gene in cancer (Vogelstein *et al*, 2000), approximately 50% of all cancers retain wild-type p53 (WT-p53) and typically circumvent or suppress p53’s functions through alternative non-mutational mechanisms (Brown *et al*, 2009). Re-awakening the latent tumour suppressive functions in cancers retaining WT-p53 is an attractive clinical concept (Brown *et al*, 2009), particularly since genetic reactivation is highly effective at inducing tumour regression in mouse models (Donehower & Lozano, 2009). Given its central importance in suppressing p53, targeting the p53-MDM2 interaction has received significant attention and has led to the development of the selective small molecule inhibitors of MDM2 (MDM2i) such as Nutlin-3A (Vassilev, 2004). While highly effective at stabilizing p53 (Brown *et al*, 2009). such agents promote variable cellular fates depending on cell type in all but *MDM2*-amplified cells, in which they induce cell death (París *et al*, 2008; Tovar *et al*, 2006). Pre-clinical and clinical development of the next generation of inhibitors of MDM2 (MDM2i) (and family member MDMX/MDM4) is ongoing (Wade *et al*, 2013), and these agents are increasingly being combined with other therapies (Zhao *et al*, 2015). The observation that potent p53 stabilisation elicited by MDM2i is often insufficient to induce cell death suggests the existence of fundamental mechanisms that regulate the cell death-inducing effects of stabilized p53. Strategies targeting such p53-mediated cell death inhibitory mechanisms have potential to not only enhance MDM2i-induced cell death, but also to augment the efficacy of p53-induced cell death induced by radiotherapy and DNA damaging chemotherapies.

Acetylation of a dense cluster of lysine residues in p53’s C-terminus, which are also target residues for ubiquitination by MDM2, has been suggested to enhance transactivation of pro-apoptotic target genes (Laptenko *et al*, 2015; Marouco *et al*, 2013; Tang *et al*, 2008a). Nuclear Class I HDACs (HDAC1/2/3) have been shown to both interact with MDM2 to enhance p53 degradation and directly deacetylate the p53 C-terminus to influence its stability and transcriptional activity (Ito *et al*, 2002; Juan *et al*, 2000; Harms & Chen, 2007; Wang *et al*, 2012; Leboeuf *et al*, 2010). HDAC activity is frequently deregulated in cancer, and cancer cells are typically more sensitive to HDAC inhibition than normal cells (Lee *et al*, 2010); hence, this class of epigenetic modifying enzymes has received significant attention as therapeutic targets, and HDAC inhibitors have achieved several clinical approvals despite rather limited understanding of their mechanisms-of-action (Guha, 2015).

In this study, we demonstrate that selective inhibition of HDAC1/2/3 using the clinically relevant agent Entinostat (Trapani *et al*, 2017) synergises with direct and indirect p53-activating agents to activate apoptotic cell death. At the heart of this synergy, we identify the apoptosis regulator FLIP’s long splice form (Fas-associated death domain (FADD)-like interleukin-1β-converting enzyme inhibitory protein) (FLIP(L)) as a direct p53 target gene and nodal determinant of cell fate following p53 activation.

## Results

### Class-I HDAC inhibition enhances p53-dependent cell death

Given the importance of acetylation (of multiple lysines in the C-terminus and DNA binding domain) of p53 in regulating cell fate (Tang *et al*, 2008b; Marouco *et al*, 2013), we investigated the impact of the pan-HDAC inhibitor Vorinostat (Suberanilohydroxamic acid, SAHA) on cell death induction in paired p53-WT and null HCT116 cells. p53-WT cells were more sensitive to apoptosis induced by SAHA compared to their p53-null counterparts, or matched cells expressing the clinically relevant DNA binding-defective mutant p53 R248W from the endogenous locus (Sur *et al*, 2009) (Supplementary information, Fig. S1a). This sensitivity correlated with enhanced acetylation of the C-terminal region of p53^K373^ (Supplementary information, Fig. S1b). Notably, despite potent effects on p53 stabilisation, the specific MDM2 inhibitor Nutlin-3A induced cell death in only 10% of p53-WT cells (Supplementary information, Fig. S1a). Importantly, addition of SAHA to cells pre-treated for 24 hours with Nutlin-3A or the clinically relevant DNA damaging agent Oxaliplatin significantly enhanced cell death in a wild-type p53-dependent manner (Fig. 1a and Supplementary information, Fig. S1c). As expected, SAHA treatment increased acetylation of HDAC targets K373 and K382 in the C-terminus of p53-WT stabilised in response to both Nutlin-3A and Oxaliplatin (Fig. 1b and Supplementary information, Fig. S1d). These effects were not observed in p53 R248W mutant knock-in cells (Supplementary information, Fig. S1a-d), indicating that enhancement of cell death requires p53 DNA-binding activity and suggesting a transcriptional mechanism mediated by inhibition of nuclear HDACs.

**Fig. 1.**
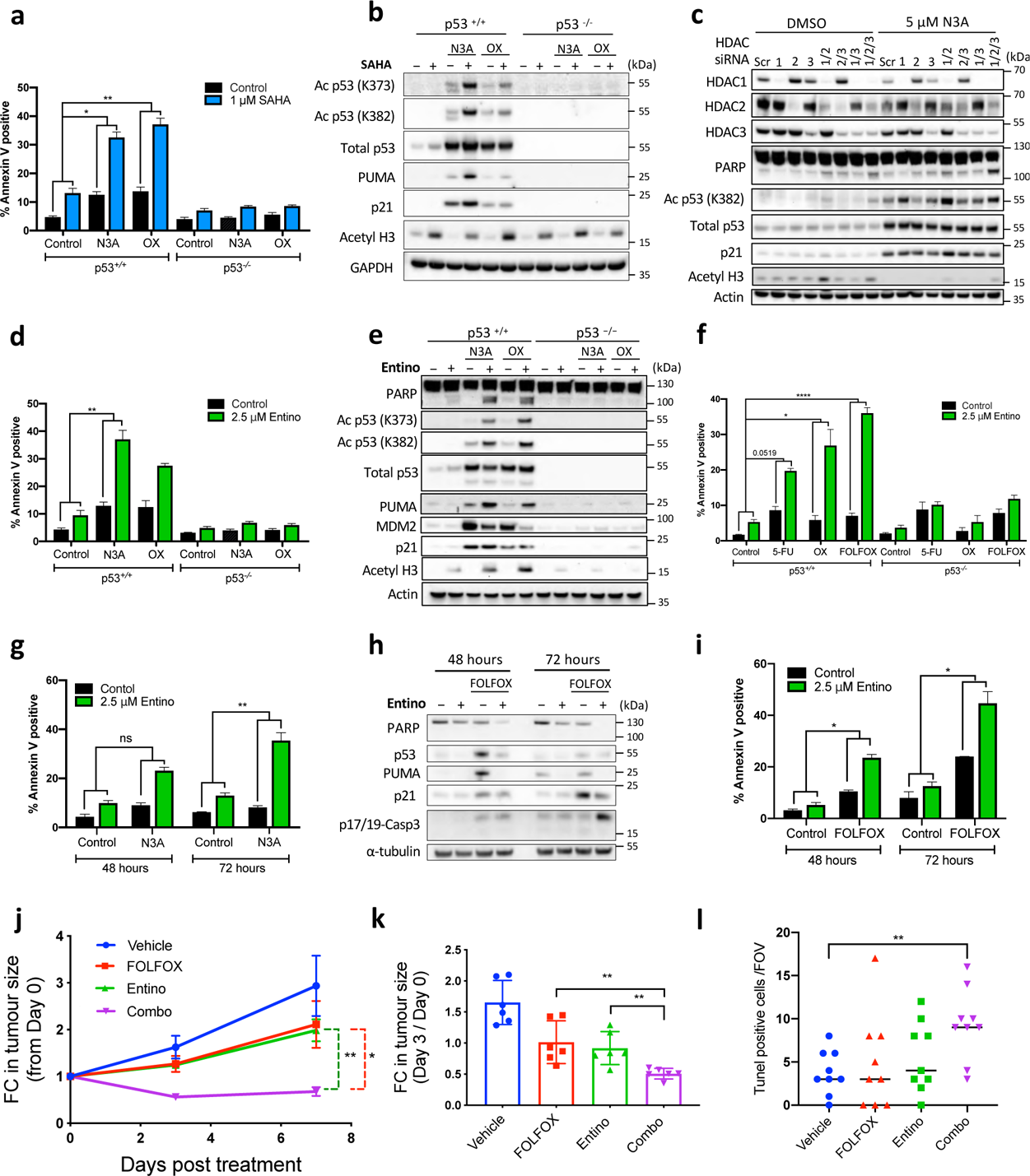
Class-I HDAC inhibition enhances p53-dependent apoptotic cell death. (**a**) Annexin-V/PI FACS assessment of cell death **(b)** and Western blot analysis of p53-WT and null HCT116 cells treated with 5 µM Nutlin-3A (N3A) or 1 µM Oxaliplatin (OX) for 24 hours (h) prior to treatment with 1 µM SAHA for a further 24 h. (**c**) Western blot analysis of protein expression in p53-WT HCT116 cells transfected with control (Scr) or HDAC1/2/3-targeted siRNAs for 24 h prior to treatment with 5 µM N3A for a further 48 h. Annexin-V/PI FACS (**d**) and Western blot (**e**) analysis of p53-WT and null HCT116 cells treated with 5 µM Nutlin-3A (N3A) or 1 µM Oxaliplatin (OX) for 24 h prior to treatment with 2.5 µM Entinostat (Entino) for a further 24 h. (**f**) Comparison of cell death induced by 2.5 µM 5-FU, 1 µM Oxaliplatin alone or their combination (FOLFOX) for 24 h prior to treatment with Entinostat (Entino) for a further 24 h in p53-WT and null HCT116 cells. (**g**) Annexin-V/PI FACS and (**h**) Western blot assessment of cell death induced after 48 h and 72 h concurrent exposure of CT26 murine CRC cells to Entinostat (2.5 µM) and FOLFOX (1.25 µM 5-FU plus 1 µM Oxaliplatin). (**i**) Annexin-V/PI FACS assessment of cell death induced following 48 h and 72 h concurrent exposure of CT26 cells to Entinostat (2.5 µM) in combination with Nutlin-3a (5 µM). (**j**) Assessment of fold change (FC) in tumour size of BALB/c mice (3 per group) injected with 1 x 10^6^ CT26 murine cells over 7 days of daily treatment with vehicle (30% cyclodextrin/PBS), FOLFOX (5-FU 10 mg/kg, Oxaliplatin 1 mg/kg), Entinostat (10 mg/kg), or their combination. (**k**) Tumour growth analysis (6 per group) of mice treated for 3 days as in (**j**). (**l**) Quantification of TUNEL-positive cells per field of view (FOV) 72 h after treatment as in (**j**). Three mice per group were analysed. Error bars in (**a/d/f/g/l**) are represented as mean ± s.e.m. of at least three independent experiments. *P* values \**P* < 0.05; \*\**P* < 0.01, \*\*\**P* < 0.001, \*\*\*\**P* < 0.0001 calculated by 2-way ANOVA. Error bars in (**j/k**) are represented as mean ± s.d. *P* values in (**j/k/l**) \**P* < 0.05; \*\**P* < 0.01, \*\*\**P* < 0.001 calculated by Students t-test.

SAHA is a broad-spectrum inhibitor, which suppresses activity of multiple HDACs, including the predominantly nuclear Class-I HDACs 1/2/3 (Bradner *et al*, 2010) responsible for acetylating p53(Juan *et al*, 2000; Leboeuf *et al*, 2010; Wang *et al*, 2012; Ito *et al*, 2002). siRNA-mediated downregulation of HDACs 1/2/3 individually or in combination revealed that simultaneous depletion of all 3 nuclear HDACs is necessary to enhance Nutlin-3A-induced apoptosis as indicated by PARP cleavage (Fig. 1c). Moreover, these effects were recapitulated when the Class-I-selective HDAC inhibitor Entinostat was combined with either Nutlin-3A or Oxaliplatin in (Fig. 1d/e). For both siRNA- and small molecule-mediated targeting of HDAC1/2/3, the cell death induced was coincident with acetylation of p53^K373/K382^ (Fig. 1c/e). Similar p53-dependent effects of Entinostat were observed in further isogenic paired CRC models (LoVo and RKO; Supplementary information, Fig. S1e-h). Combining Entinostat with the backbone chemotherapeutic agent 5-Fluorouracil (5-FU) and the clinically relevant chemotherapy doublet of 5-FU and Oxaliplatin (“FOLFOX”) also resulted in p53-dependent enhancement of cell death in HCT116 cells (Fig. 1f). Similar effects on cell death were observed in response to combination of Entinostat with FOLFOX or Nutlin-3A in the p53 wild-type CT26 murine model of microsatellite stable (MSS), *KRAS* mutant CRC (Fig. 1g-i). Importantly, Entinostat significantly enhanced the *in vivo* efficacy of FOLFOX in CT26 syngeneic allografts (Fig. 1j/k), and the combined treatment was well-tolerated. Moreover, the tumour regressions observed in response to the combination treatment correlated with enhanced TUNEL positivity (Fig. 1l), indicative of an apoptotic mechanism of action.

### Entinostat modulates transcription of a subset of p53 target genes

To further assess the importance of p53-mediated transcription for the observed effects on cell death, we next evaluated the Nutlin-3A/Entinostat combination in HCT116 p53-null cells retrovirally reconstituted to express either the clinically relevant R248W DNA binding p53 mutant, a multiple Lys-Arg (8KR) acetylation mutant (Drosten *et al*, 2014) or p53-WT. Significant combinatorial effects of Nutlin-3A/Entinostat on cell death were detected only in the p53-null cells re-constituted with p53-WT (Fig. 2a/b), indicating that both DNA-binding *and* acetylation of p53 are required for the observed cell death phenotype. Therefore, we next utilized mRNA-seq to investigate the transcriptional changes underpinning this synergy. Samples from HCT116 p53-WT and p53-null cells were collected 8 hours following treatment with Nutlin-3A, Entinostat or their combination, a timepoint at which p53 was stabilised and significant increases in expression of a range of direct p53 targets (including p21/*CDKN1A*, TRAIL-R2/*TNFRSF10B* and PUMA/*BBC3*) as observed by qRT-PCR; importantly however, as transcription may be indirectly suppressed in apoptotic cells, this 8-hour timepoint precedes activation of apoptotic cell death (Supplementary information, Fig. S2A-B).

**Fig. 2.**
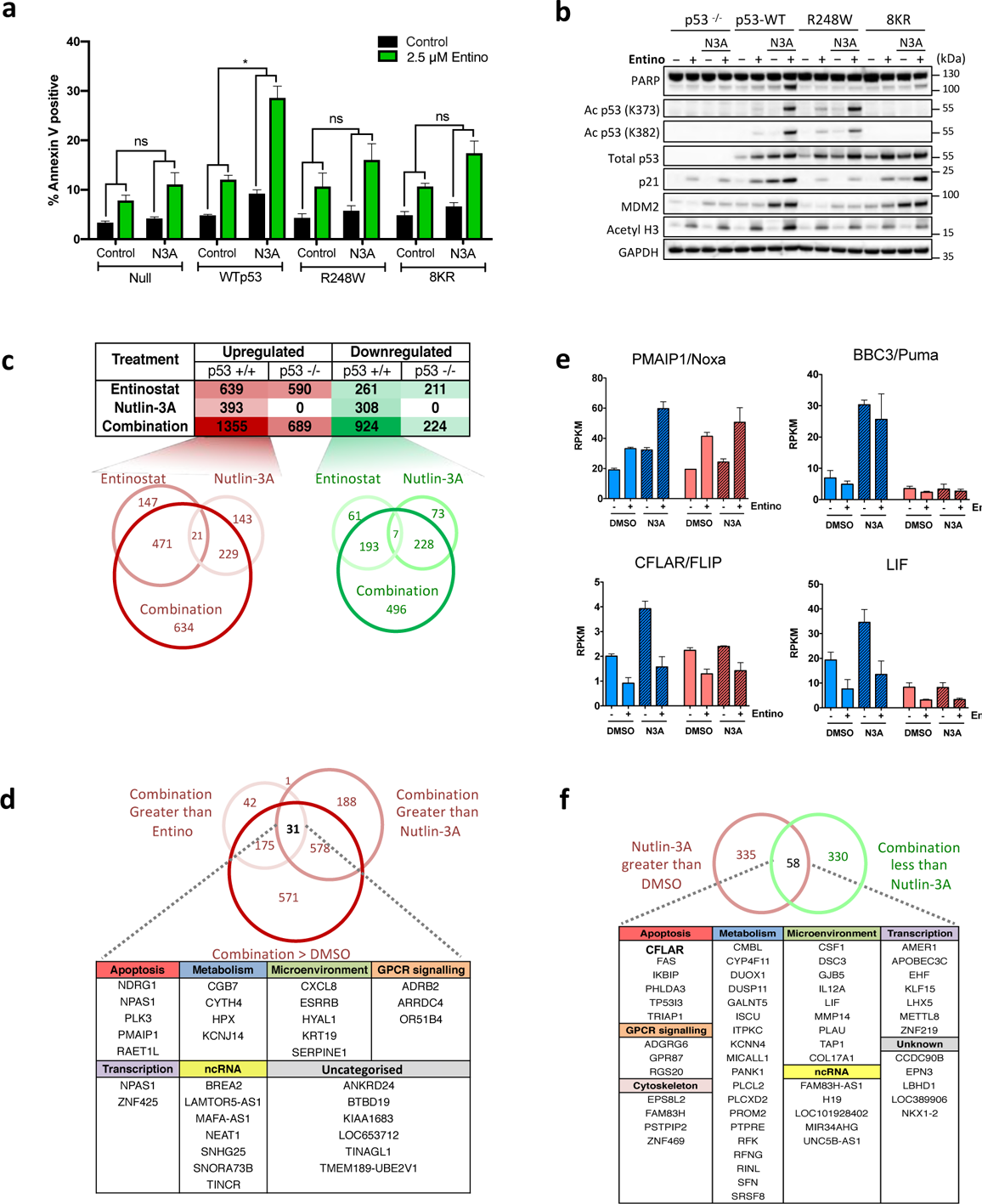
Entinostat modulates transcription of a subset of p53 target genes. (**a**) Annexin-V/PI FACS assessment of cell death and (**b**) Western blot analysis of parental p53-null HCT116 cells and daughter cells ectopically expressing wild-type (WTp53), R248W mutant or 8KR acetylation mutant p53. Cells were treated with 5 µM Nutlin-3A (N3A) for 24 hours (h) prior to treatment with 2.5 µM Entinostat (Entino) for an additional 24 h. (**c**) Differential gene expression analysis of mRNA-seq performed in HCT116 p53 wild-type (WT) and null cells from two independent experiments. RNA was extracted 8 h post treatment with DMSO, 5 µM Nutlin-3A, 2.5 µM Entinostat or their combination. Mapped read counts for transcripts were RPKM normalized for relative expression, and differentially expressed transcripts between each treatment detected by 3-way ANOVA (mean RPKM ≥1.7x, FDR 0.05). The total numbers of significantly up/downregulated genes vs DMSO for each treatment are tabulated. Venn diagrams illustrating numbers of transcripts significantly upregulated (Red) or downregulated (Green) in response to Entinostat, Nutlin-3A and the combination in p53-WT cells. (**d**) Venn diagram of numbers of transcripts significantly upregulated in response to the combination treatment compared to both single agent treatments in p53-WT cells, and list of 31 cooperatively upregulated transcripts, sub-categorized based on cellular function. (**e**) RPKM values for functionally-related p53 targets *PMAIP1*/Noxa (from panel **d** and **f**) and *BBC3*/PUMA represented as mean RPKM ± s.d. of two independent experiments. (**f**) Venn diagram of numbers of transcripts whose transcriptional upregulation in response to Nutlin-3A is significantly attenuated in the combination treatment, and list of those 58 genes sub-categorized based on cellular function. Error bars in (**a**) are represented as mean ± s.e.m. of at least three independent experiments. *P* values \**P* < 0.05; \*\**P* < 0.01, \*\*\**P* < 0.001 calculated by 2-way ANOVA.

Treatment of p53-WT HCT116 cells with Nutlin-3A alone resulted in a significant increase (≥1.7-fold) in 393 genes (Fig. 2c; Supplementary information, Table S1), as expected, enriched for p53 signalling, apoptosis and direct p53 target genes (Supplementary information, Table S2a). Even at this early time-point, 308 significantly repressed genes were identified; these were enriched for cell cycle and FOXM1/E2F4 targets potentially mediated through indirect suppression downstream of p21 activation (reviewed in (Fischer *et al*, 2014)) (Supplementary information, Table S1 and Supplementary information, Table S2a). No significantly altered genes were identified in the p53-null model in response to Nutlin-3A (Fig. 2c), underlining the selectivity of this MDM2 inhibitor. Conversely, there was significant overlap between the genes induced or repressed by Entinostat in both models, indicating that a large proportion of the effects of single agent Entinostat at this timepoint were not affected by p53 status (Supplementary information, Table S1 and Supplementary information, Table S2b); most notable was p53-independent suppression of NFκB.

When we further compared the effects of the Nutlin-3A/Entinostat combination in p53-WT cells, only 31 genes were identified to be significantly upregulated relative to single agent treatments (Fig. 2d and Supplementary information, Table S1/2d). It was notable that only one of these, Noxa/*PMAIP1*, was a canonical direct p53 induced pro-apoptotic target (Fig. 2c-e, and confirmed by qRT-PCR in Supplementary information, Fig. S2b). Surprisingly, 143 Nutlin-3A-induced genes were not significantly upregulated in combination-treated cells (Fig. 2c and Supplementary information, Table S2c), of which 58 were significantly reduced compared to Nutlin-3A treatment alone (Fig. 2f and Supplementary information, Tables S1/2). Interestingly, both gene lists were significantly enriched for tumour necrosis factor (TNF) and p53 signalling pathways (Fig. 2f and Supplementary information, Table S2d); this list included both pro- and anti-apoptotic regulators of death receptor signalling, for example the pro-apoptotic death receptor p53 target *FAS*/FAS (a canonical p53 target) and the regulator of death receptor-induced, caspase-8-mediated apoptosis *CFLAR*/FLIP (Fig. 2e/F and Supplementary information, Fig. S2b/c).

### Functional genomics analyses identify *CFLAR*/FLIP as a direct HDAC1/2/3-dependent pro-survival target of p53

To investigate the mechanisms underlying Entinostat-enhanced, p53-dependent cell-death, we evaluated the effect of Nutlin-3A in HCT116 cells transfected with a panel of siRNAs targeting 37 core cell death regulators (Fig. 3a). Aligning these results with the mRNA-seq dataset identified *CFLAR*/FLIP as a potential nodal regulator of p53-induced death since its depletion significantly enhanced the effects of Nutlin-3A on viability and its mRNA expression was induced in a p53-dependent manner, but attenuated by Entinostat co-treatment (Fig. 2e, 3a and Supplementary information, Table S1/2). Moreover, we also identified sgRNAs targeting *CFLAR*/FLIP as one of the top sensitizers to Nutlin-3A in gene-level MAGeCK analysis of a genome-wide CRISPR screen (Fig. 3b and Supplementary information, Table S3) further supporting a nodal role for *CFLAR*/FLIP in promoting survival in cells in which p53 is activated. As a quality control for the screen, sgRNAs targeting p53 and *CDKN1A*/p21 were identified as mediators of resistance to Nutlin-3A (Supplementary information, Fig. S3a). Re-analysis of previous ChIP-seq data from our group in human foreskin keratinocytes treated with cisplatin (McDade *et al*, 2014) or public MCF7 breast cancer cells treated with Nutlin-3A (Nikulenkov *et al*, 2012) identified direct, treatment-induced binding of p53 to the *CFLAR*/FLIP promoter (Supplementary information, Fig. S3b), and this was confirmed in HCT116 cells following treatment with Nutlin-3A (Fig. 3c). This coincided with induction of transcription of both the long FLIP(L) and short splice forms FLIP(S) (the latter expressed at significantly lower levels) as detected by qRT-PCR (Fig. 3d). This analysis also confirmed that Nutlin-3A-induced CFLAR/FLIP mRNA upregulation was attenuated by co-treatment with Entinostat, with similar effects observed in the LoVo model (Supplementary information, Fig. S2c).

**Fig. 3.**
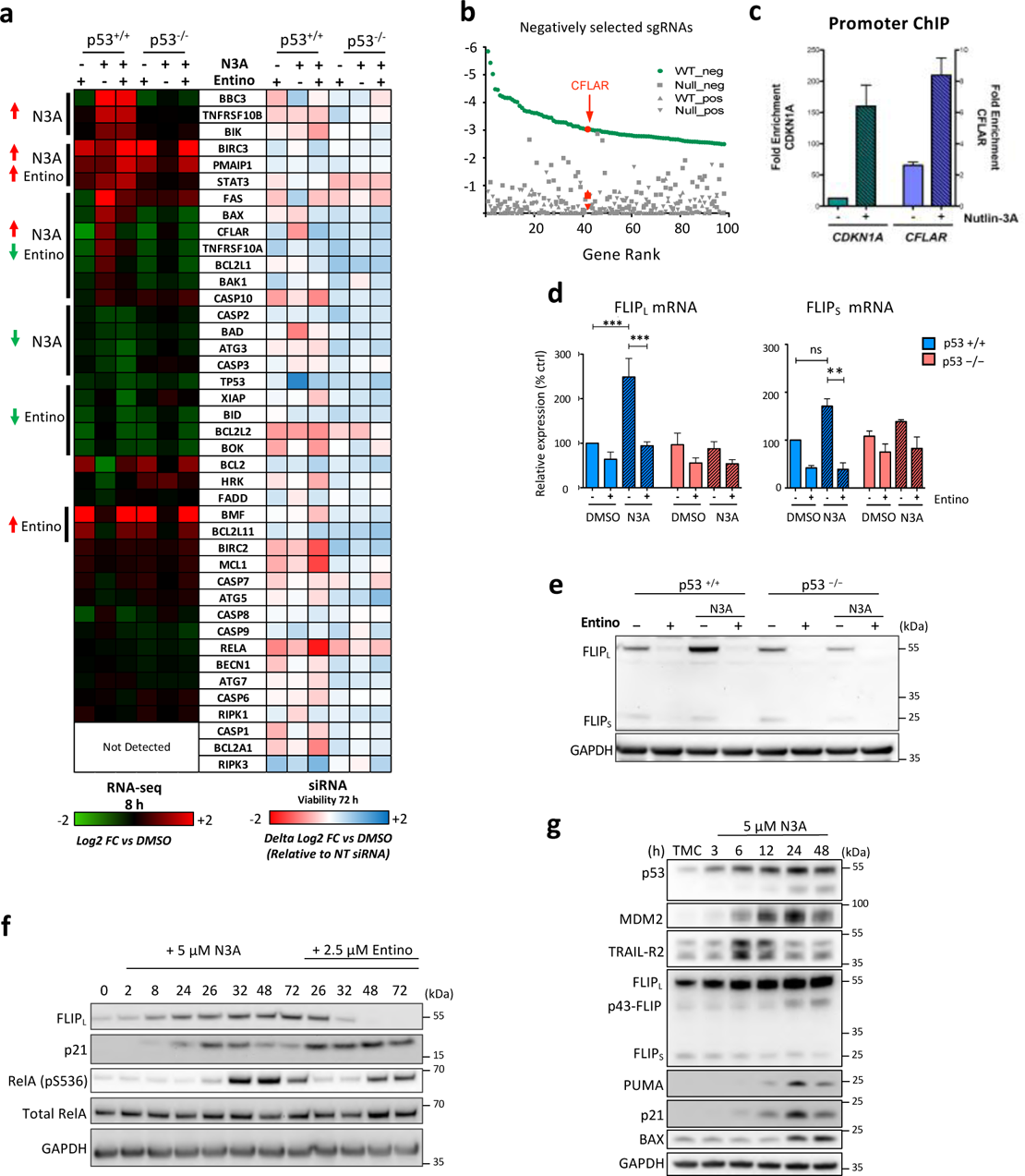
Functional genomics analyses identify *CFLAR*/FLIP as a direct HDAC1/2/3-dependent pro-survival target of p53. (**a**) (*Left side*) Heatmap of RNA-seq data of a panel of apoptosis-regulatory genes from p53-WT and null HCT116 cells treated with 2.5 µM Entinostat (Entino), 5 µM Nutlin-3A (N3A) or their combination for 8 h. Mean RPKM fold-changes from 2 independent experiments are log2-transformed, and associated colour key contiguous with log2 expression values vs DMSO. (*Right side*) Cell viability data from p53-WT and null HCT116 cells transfected with the indicated siRNAs for 24 h, followed by treatment with 5 µM Nutlin-3A for 72 h. Values represent delta log2 fold change of Nutlin-3A vs DMSO derived from average log2 fold changes in cell viability vs non-targeting (NT) siRNA from 2 independent experiments. No response to Nutlin-3A is displayed as white, with subsequent increases or decreases in sensitivity represented as red and blue respectively. (**b**) Results of a genome-wide CRISPR screen in HCT116 p53-WT or Null cells for enhancers of Nutlin-3A sensitivity or resistance. The *p*-value for MAGeCK gene level analysis of top 100 genes whose CRISPR guide RNAs (sgRNAs are negatively (sensitisers) or positively (resistance) selected CRISPR guide RNAs (sgRNAs) are depicted, highlighting p53 (*TP53*) and p21 (*CDKN1A*) and FLIP (*CFLAR*). (**c**) p53 Chromatin Immunoprecipitation (ChIP) assay in p53-WT HCT116 cells using primers directed against the FLIP/*CFLAR* promoter region. Relative promoter occupancy between treatments was calculated by fold-enrichment of target region versus a non-specific region (Cyclin D1/*CCND1*). Data are represented as mean ± s.e.m of two independent experiments. (**d**) Quantitative RT-PCR of FLIPL and FLIPS mRNA expression. Data are normalized to RPL24 control for each sample. (**e**) Western blot assessment of FLIPL and FLIPS protein expression in p53-WT and null HCT116 cells treated for 24 h with 5 µM Nutlin-3A (N3A) with or without 2.5 µM Entinostat (Entino). (**f**) Western blot analysis of p53-WT HCT116 cells treated with 5 µM N3A with or without 2.5 µM Entino for the indicated times. (**g**) Western blot analysis of HCT116 cells treated with 5 µM N3A for the indicated times (TMC = time matched control). Error bars in (**d**) are represented as mean ± s.e.m. of at least three independent experiments. *P* values \**P* < 0.05; \*\**P* < 0.01, \*\*\**P* < 0.001 calculated by 1-way ANOVA.

Changes in CFLAR/FLIP mRNA expression in HCT116 p53-WT cells were reflected at the protein level for FLIP(L) but not for FLIP(S), which was expressed at a much lower level than the long splice form (Fig. 3e and Supplementary information, Fig. S3c). Since *CFLAR*/FLIP is a known NFκB target gene (reviewed in (Riley *et al*, 2015)) and our mRNA-seq analysis indicated the potential for Entinostat to significantly suppress components of the NFκB pathway in both p53-WT and -null cells (Supplementary information, Table S2b), we next evaluated whether NFκB contributed to changes in FLIP expression. Time-course analysis in both HCT116 and LoVo cells indicated that FLIP(L) protein expression was induced at early time-points coincident with or even preceding p21 upregulation (Fig. 3f and Supplementary information, Fig. S3d/e), whereas activation of the canonical NFκB pathway as assessed by phospho-RelA was not observed until later time-points (Fig. 3f and Supplementary information, Fig. S3e). In addition, while the absolute levels of FLIP(L) expression were decreased by the IKK inhibitor BAY-117082 (confirming FLIP/*CFLAR* as an NFκB regulated gene), the magnitude of p53-dependent induction of FLIP(L) was maintained (Supplementary information, Fig. S3g), with similar results obtained in HCT116 stably transduced with the IκB-super-repressor (Luo *et al*, 2004) (Supplementary information, Fig. S3f). Upregulation of FLIP(L) coincided with TRAIL-R2 upregulation, both of which preceded upregulation of the canonical p53 apoptotic targets BAX and PUMA (Fig. 3g). At later timepoints, the FLIP(L) cleavage product p43-FLIP was also detectable (Fig. 3g); when FLIP(L) forms FADD-dependent heterodimers with procaspase-8, it is cleaved in its C-terminus between its large and small pseudo-catalytic domain(Humphreys *et al*, 2018). Altogether, these results establish FLIP and particularly FLIP(L) as a direct p53 target that is induced in an independent manner that is independent from its canonical regulation by NFκB.

### Nutlin-3A induces FLIP(L)-dependence by priming p53-WT CRC cells for caspase-8-dependent apoptosis

To investigate the functional importance of Nutlin-3A-induced dependence on FLIP for suppressing p53-induced cell death, we first confirmed the predictions of the siRNA and CRISPR screens by demonstrating that RNAi-mediated depletion of FLIP synergistically enhances cell death induced by Nutlin-3A treatment in a p53-dependent manner (Fig. 4a/b). These results were further confirmed in LoVo and RKO p53-WT CRC cell line models (Supplementary information, Fig. S4a/b). Moreover, splice form-specific siRNA indicated that in HCT116 cells selective FLIP(L) depletion was sufficient to promote Nutlin-3A-mediated cell death, with depletion of FLIP(S) (which is not induced by p53 at the protein level) having only a modest effect on PARP cleavage (Supplementary information, Fig. S4c).

**Fig. 4:**
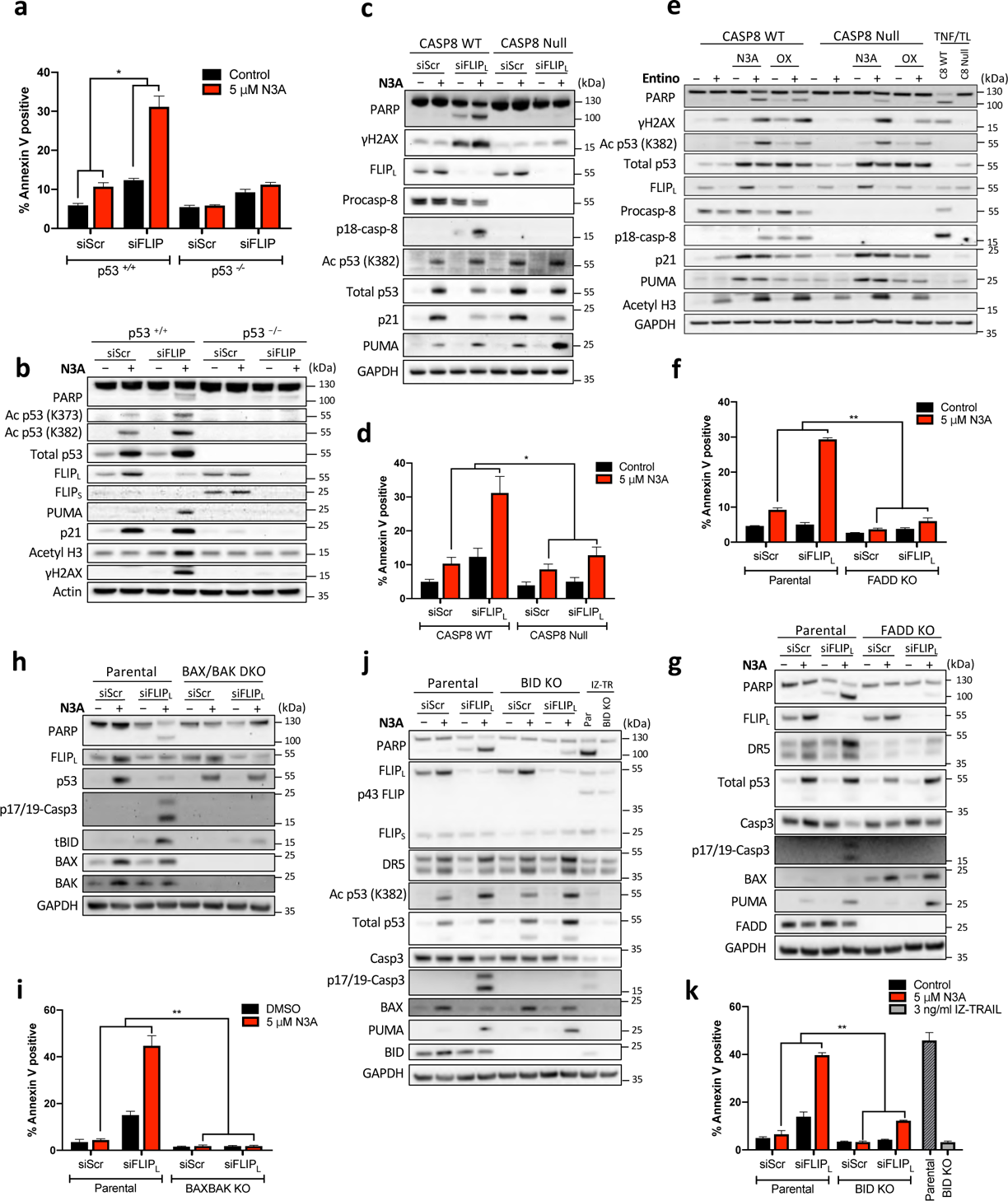
Nutlin-3A induces FLIP(L)-dependence by priming p53 wild-type CRC cells for caspase-8-dependent apoptosis. (**a**) Annexin-V/PI FACS and (**b**) Western blot assessment of cell death of HCT116 p53-WT and null cells transfected with 10 nM FLIP or control siRNA for 6 h prior to treatment with 5 µM Nutlin-3a (N3A) for a further 24 h. (**c/g/h/j**) Western blot and (**d/f/i/k)** Annexin-V/PI FACS analysis of effects of siRNA mediated depletion of FLIPL in combination with 5 µM N3A as in panel (A/B) in HCT116 isogenic knockout models for caspase-8 (CASP8) (C/D), FADD (F/G), BAX/BAK DKO (**h/i**) and BID (**j/k**) (treatment with 3 ng/ml isoleucine zipper TRAIL (IZ-TRAIL) serves as a positive control). (**e**) Western blot assessment of caspase 8 WT and null cells pre-treated for 24 h with either 5 µM N3A or 1 µM Oxaliplatin (OX) prior to treatment of 2.5 µM Entinostat for a further 24 h. 24 h treatment with 10 ng/ml TNFɑ/1 µM Biranapant (TL) serves as a positive control. Error bars in (**a/d/f/i/k**) are represented as mean ± s.e.m. of at least three independent experiments. *P* values \**P* < 0.05; \*\**P* < 0.01; *** < 0.001 calculated by 2-way ANOVA.

The best described function of FLIP is its role as a regulator of caspase-8-dependent apoptosis; however, whether FLIP(L) acts an inhibitor or activator of caspase-8 remains a matter of debate. We therefore extended our analyses to a CRISPR-Cas9 caspase-8-deficient HCT116 model (Paek *et al*, 2016), which was found to be significantly more resistant to the cell death inducing effects of FLIP(L) silencing in cells treated with Nutlin-3A for 24 h than control caspase-8-expressing cells (Fig. 4c/d). Similarly, the apoptosis induced by Entinostat in combination with Nutlin-3A or oxaliplatin was also significantly attenuated in cells lacking caspase-8 (Fig. 4e). FLIP and caspase-8 interact via the adaptor protein FADD (Humphreys *et al*, 2018), and the cell death induced by Nutlin-3A/siFLIP(L) was also FADD-dependent (Fig. 4f/g). HCT116 cells have been identified by us and others as “Type-II” in terms of caspase-8-mediated apoptosis (Wilson *et al*, 2008) meaning that they require amplification of extrinsically-derived apoptotic signals via the mitochondria. Consistent with this, the apoptosis induced by Nutlin-3A/siFLIP(L) was completely attenuated in BAX/BAK double knockout (DKO) cells (Fig. 4h/i) and also partially attenuated in SMAC null cells (Supplementary information, Fig. S4d/e). In the extrinsic pathway, crosstalk between caspase-8 and the mitochondria is mediated by the BH3 protein BID, which when cleaved by caspase-8 to its truncated form (tBID) translocates to the mitochondria and promotes mitochondrial outer membrane permeabilization (MOMP). Consistent with this, the apoptosis induced by Nutlin-3A/siFLIP(L) was significantly attenuated, although not completely prevented, in BID CRISPR knock-out cells (Fig. 4j/k). By comparison, the effects of the direct extrinsic apoptosis pathway activator isoleucine zipper-TRAIL (IZ-TRAIL) were completely abrogated by BID deletion, confirming this model as “Type 2” (Fig. 4j/k).

### Nutlin-3A induces formation of a ligand-independent TRAIL-R2 complex

FLIP and FADD regulate procaspase-8 processing and activation at death-inducing signaling complexes (DISCs) formed by death receptors and the cytoplasmic ripoptosome formed by RIPK1 (Riley *et al*, 2015). To identify the upstream regulator(s) of caspase-8-mediated apoptosis in Nutlin-3A-treated, FLIP(L)-depleted cells, we re-examined the mRNA-Seq data for the effects of Nutlin-3A and Entinostat on death receptor expression. *FAS*/Fas and *TNFRSF10A*/DR4/TRAIL-R1 were both upregulated in response to Nutlin-3A, but this increase was suppressed by co-treatment with Entinostat (data viewable in HDAC_visualiseR Shiny App). *TNFRSF10B/*DR5/TRAIL-R2, a known direct p53 target (Takimoto & el-Deiry, 2000), was expressed at a higher level than the other 2 death receptors and was as expected significantly upregulated in response to Nutlin-3A; however in this case, the upregulation was unaffected by co-treatment with Entinostat. RIPK1 was expressed at a low level that was unaffected transcriptionally by either treatment. Notably, RNAi-mediated downregulation of TRAIL-R2, but not the other death receptors nor RIPK1 (*data not shown*), significantly rescued the cell death induced by treatment with Nutlin-3A in combination with siFLIP(L) and did so to a similar extent as silencing caspase-8 and p53 (Fig. 5A). Of note, cleavage and activation of BID was also reduced in TRAIL-R2- and caspase-8-depleted cells (Fig. 5a). These results were confirmed in 2 independent p53-WT CRC models (Supplementary information, Fig. S5a) and using an HCT116 model in which TRAIL-R2 expression was deleted by CRISPR-cas9 (Fig. 5b/c). Silencing TRAIL-R2 or CASP8 (and as expected p53) in HCT116 cells also partially inhibited apoptosis signaling in response to Nutlin-3A/Entinostat co-treatment (Fig. 5d).

**Fig. 5.**
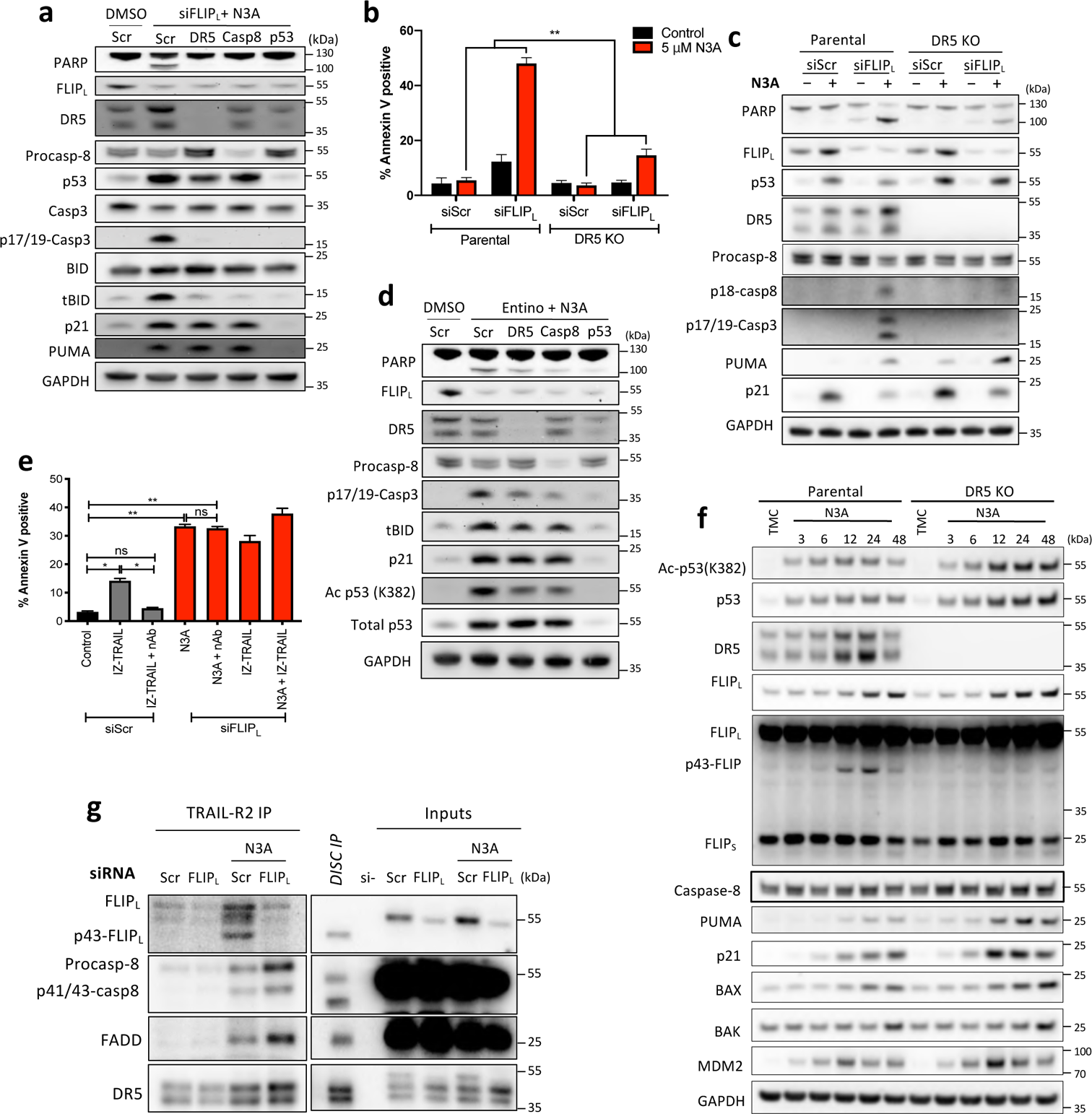
Nutlin-3A induces formation of a ligand-independent TRAIL-R2 complex. (**a**) Western blot assessment of HCT116 p53+/+ cells transfected with the indicated siRNA’s (10 nM) for 24 h followed by transfection with siFLIPL for 6 h and treatment with 5 µM Nutlin-3A (N3A) for a further 24 h. (**b**) Annexin-V/PI FACS and (**c**) Western blot assessment of cell death in HCT116 DR5/TRAIL-R2 CRISPR KO and matched parental cells transfected for 6 h with 10 nM scrambled (siScr) or a FLIPL siRNA prior to treatment with 5 µM Nutlin-3A (N3A) for a further 24 h. (d) Western blot assessment of HCT116 p53+/+ cells transfected with the indicated siRNA’s (10 nM) for 24 h followed by co-treatment with 5 µM Nutlin-3A (N3A) and 2.5 µM Entinostat (Entino) for 24 h. (**e**) Annexin-V/PI FACS analysis of the effect of TRAIL neutralizing antibody treatment (100 ng/ml) on cell death in cells transfected for 6 h with 10 nM scrambled (siScr) or FLIPL siRNA prior to treatment with 3 ng/ml isoleucine zipper TRAIL (IZ-TRAIL) and/or 5 µM Nutlin-3A (N3A) for a further 24 h. (**f**) Western blot assessment of HCT116 DR5/TRAIL-R2 CRISPR KO and matched parental cells treated with 5 µM Nutlin-3A (N3A) for the indicated times (TMC = time matched control). (**g**) Post-lysis DR5/TRAIL-R2 IP performed in HCT116 parental cells transfected for 6 h with 10 nM scrambled (siScr) or a FLIPL siRNA prior to treatment with 5 µM Nutlin-3A (N3A) for a further 24 h. Error bars in (**b/e**) are represented as mean ± s.e.m. of at least three independent experiments. *P* values in (c) \**P* < 0.05; \*\**P* < 0.01; *** < 0.001 calculated by 2-way ANOVA. *P* values in (**e**) \**P* < 0.05; \*\**P* < 0.01; *** < 0.001 calculated by 1-way ANOVA.

Canonically, TRAIL-R2-mediated activation of caspase-8 and apoptosis involves binding of its ligand TRAIL (Humphreys *et al*, 2018); it was therefore highly notable that co-incubation with a TRAIL neutralizing antibody failed to rescue cell death induced by Nutlin-3A/siFLIP(L) (Fig. 5e), a result confirmed in both RKO and LoVo models (Supplementary information, Fig. S5b/c). Moreover, addition of IZ-TRAIL did not significantly enhance cell death induced by Nutlin-3A/siFLIP(L) (Fig. 5e and Supplementary information, Fig. S5d). As expected, FADD-, CASP8/10-, BID- and BAX/BAK-knockout models were resistant to IZ-TRAIL; however, the TRAIL-R2-knockout model (which still expresses TRAIL-R1/DR4; *data not shown*) retained IZ-TRAIL sensitivity (Supplementary information, Fig. S5e/f); this is further evidence that TRAIL-R2-dependent, Nutlin-3A/siFLIP(L)-induced apoptosis operates via a mechanism that is distinct from cell surface ligation of the receptor. Furthermore, in time-course experiments, the upregulation of TRAIL-R2 and FLIP(L) coincided with the generation of the p43-form of FLIP(L), which is normally only generated at the DISC, and notably, the generation of p43-FLIP(L) was almost completely attenuated in TRAIL-R2 knockout cells (Fig. 5f).

Collectively, these results suggested formation of a novel ligand-independent TRAIL-R2/FLIP(L) complex in cells treated with Nutlin-3A. To assess this, we isolated TRAIL-R2 complexes formed in response to Nutlin-3A (in the absence of ligand) using the humanized anti-TRAIL-R2 antibody Conatumamab (AMG655). This analysis indicated that Nutlin-3A indeed stimulates formation of a complex between TRAIL-R2 and FADD, FLIP(L) and caspase-8 (Fig. 5g). Importantly, both FLIP(L) and caspase-8 are processed in this ligand-independent complex (indicative of formation of catalytically active dimers) and depletion of FLIP(L) using siRNA enhanced caspase-8 recruitment (Fig. 5g), consistent with the apoptotic phenotype observed in FLIP(L)-depleted, Nutlin-3A-treated cells. Thus, acute p53-mediated upregulation of FLIP(L) prevents caspase-8-dependent apoptosis induction mediated in a TRAIL-independent manner by TRAIL-R2, which is concomitantly upregulated in response to Nutlin-3A.

### Later cell death is induced by Nutlin-3A/siFLIPL in the absence of caspase-8

At later timepoints (48 hours), the apoptotic cell death induced by Nutlin-3A/siFLIP(L) was notably less caspase-8-dependent than at 24 hours (Fig. 6a). Moreover, processing of procaspase-10, a paralog of procaspase-8 and FLIP, was observed in both *CASP8-*WT and -null cells in response to Nutlin-3A/siFLIP(L) (Fig. 6a). Similar to FLIP(L), the role of caspase-10 as a promoter or inhibitor of apoptosis has been controversial (Horn *et al*, 2017a); we therefore investigated whether caspase-10 could compensate for the loss of caspase-8 at later time-points. In siRNA depletion experiments, siCASP10 partially rescued PARP cleavage induced by Nutlin-3A/siFLIP(L) in caspase-8-deficent cells and did so to a similar extent as siTRAIL-R2 (Fig. 6b), indicating that this later cell death is also partly TRAIL-R2-dependent. To confirm these results, we generated caspase-10 knockout models in caspase-8-proficient and -deficient backgrounds using CRISPR-Cas9 (Fig. 6c/d). In the Caspase-8 wild-type setting, loss of caspase-10 did not significantly reduce the levels of apoptosis induced by Nutlin-3A/siFLIP(L) (Fig. 66c/d). However, in caspase-8-WT cells, loss of caspase-10 further decreased the cell death induced by Nutlin-3A/siFLIP(L) at 48 hours (Fig. 6c/d). Caspase-10 knockout in caspase-8-deficient cells also further inhibited apoptosis induced by Nutlin-3A/Entinostat compared to knockout of caspase-8 alone, although this was only evident at 24 hours, as by 48 hours co-deletion of caspase-8/10 failed to rescue the cell death induced by the Nutlin-3A/Entinostat combination (Supplementary information, Fig. S6a-d). Collectively, these results show that in the absence of caspase-8, caspase-10 can promote apoptosis induced by Nutlin-3A in FLIP(L)-depleted cells. In contrast, caspase-10 was unable to compensate for loss of caspase-8 to activate IZ-TRAIL-induced apoptosis (Supplementary information, Fig. S4f).

**Fig. 6.**
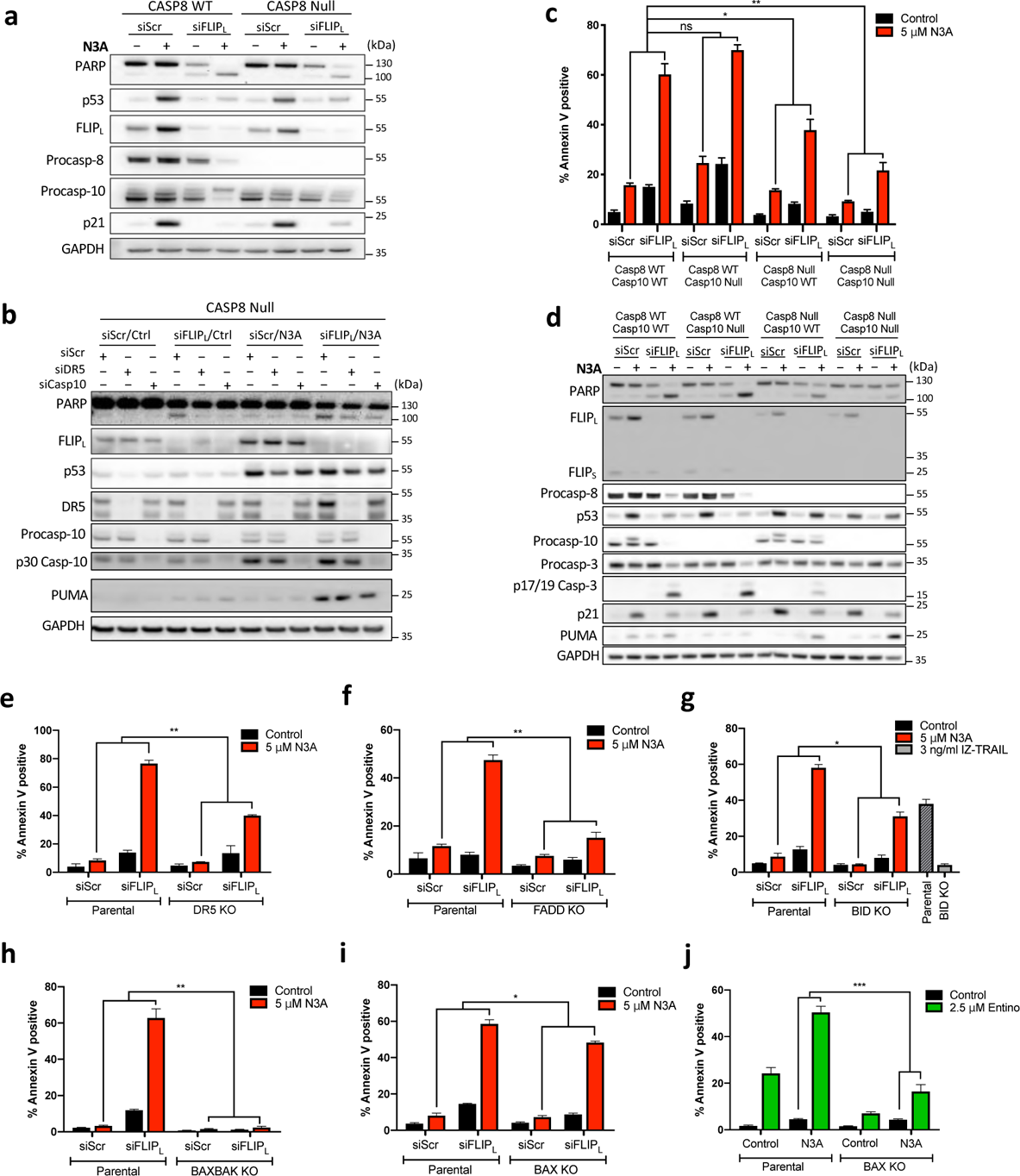
Late Cell death is induced by Nutlin-3A/siFLIPL in the absence of caspase-8. (**a**) Western blot analysis of HCT116 Caspase-8 Parental and CRISPR KO cells transfected with 10 nM FLIPL or control siRNA for 6 h prior to treatment with 5 µM N3A for a further 48 h. (**b**) Western blot analysis of HCT116 Caspase-8 CRISPR KO cells transfected with siRNA targeting DR5, Caspase-10 or their combination for 24 h prior to subsequent transfection of 10 nM FLIPL specific siRNA prior to treatment with 5 µM N3A for a further 48 h. (**c**) Annexin-V/PI FACS and (d) Western blot analysis of effects of FLIPL siRNA/N3A treatment as in (**a**) in HCT116 Caspase-8/10 CRISPR single and dual KO cells. (**e-i**) Annexin-V/PI FACS analysis of cell death in HCT116 isogeneic for DR5/TRAIL-R2 (**e**), FADD (**f**), BID (**g**), BAX/BAK DKO (**h**) and BAX (**i**) knockout treated with FLIPL siRNA/N3A as in (**a/c**). (**j**) Annexin-V/PI FACS analysis of cell death in HCT116 isogenic for BAX knockout treated with 2.5 µM Entinostat (Entino) in combination with 5 µM N3A. Error bars in (**c/e-j**) are represented as mean ± s.e.m. of at least three independent experiments. *P* values \**P* < 0.05; \*\**P* < 0.01; *** < 0.001 calculated by 2-way ANOVA.

Surprisingly, even in the absence of both caspases-8 and −10, the combination of Nutlin-3A and siFLIP(L) still induced significant levels of apoptosis at 48 hours compared to Nutlin-3A alone (Fig. 6c/d). TRAIL-R2, FADD and BID KO also significantly reduced cell death at 48 hours, however like caspase-8/10 double knockout model, significant levels of cell death were also observed when compared to Nutlin-3A treatment alone (Fig. 6e-g and Supplementary information, Fig. S6E-G). Collectively, this implies that FLIP(L) has functions that are independent of its canonical interactome. The cell death induced by Nutlin-3A/siFLIP(L) at the later timepoint was however completely attenuated in cells lacking BAX and BAK even though BID was cleaved in this model (Fig. 6h and Supplementary information, Fig. S7a), indicating involvement of the intrinsic pathway. Knockout of both BAX and BAK (and as expected p53) also completely rescued the cell death induced by Nutlin-3A/Entinostat (Supplementary information, Fig. S7b-e). Notably however, the single deletion of BAX conferred greater resistance to the apoptotic cell death induced by Nutlin-3A/Entinostat than Nutlin-3A/siFLIP(L) (Fig. 6i/j and Supplementary information, Fig. S7f/g). These results indicate overlapping but non-identical mechanisms-of-action for siFLIP(L) and Entinostat in combination with Nutlin-3A.

### FLIP(L) inhibits p53-mediated upregulation of PUMA

It was notable that Nutlin-3A-induced upregulation of the canonical p53 target PUMA (which we had initially expected to be enhanced by HDAC inhibition) was consistently enhanced in FLIP(L)-depleted cells (Fig. 4b/c/g/j, 5d, 6b/d); this was particularly apparent in cell lines in which apoptosis induction was inhibited. We therefore assessed whether PUMA was responsible for mediating the cell death phenotype induced at later timepoints by Nutlin-3A/siFLIP(L) in cells lacking caspase-8/10. In caspase-8/10-deficient models, PUMA was again found to be upregulated in response to Nutlin-3A/siFLIP(L) (Fig. 7a-c). Co-silencing of PUMA indeed rescued the residual cell death induced by Nutlin-3A/siFLIP(L) in caspase-8/10-deficient models, but had no effect in caspase-8- and/or caspase-10-proficient cells (Fig. 7a-c). Further analysis of mRNA expression indicated that silencing FLIP(L) enhanced Nutlin-3A-induced upregulation of PUMA mRNA (Fig. 7d), which notably was concomitant with suppression of CDKN1A/p21 mRNA induction, consistent with observations made at the protein level (Fig. 7b, 6a/d, 5d). Collectively, these results indicate that p53-mediated upregulation of FLIP(L) suppresses p53-mediated upregulation of PUMA and promotes p53-mediated upregulation of p21, suggestive of a novel negative feedback from FLIP(L) to suppress PUMA-mediated apoptosis and promote p21-mediated cell cycle arrest.

**Fig. 7.**
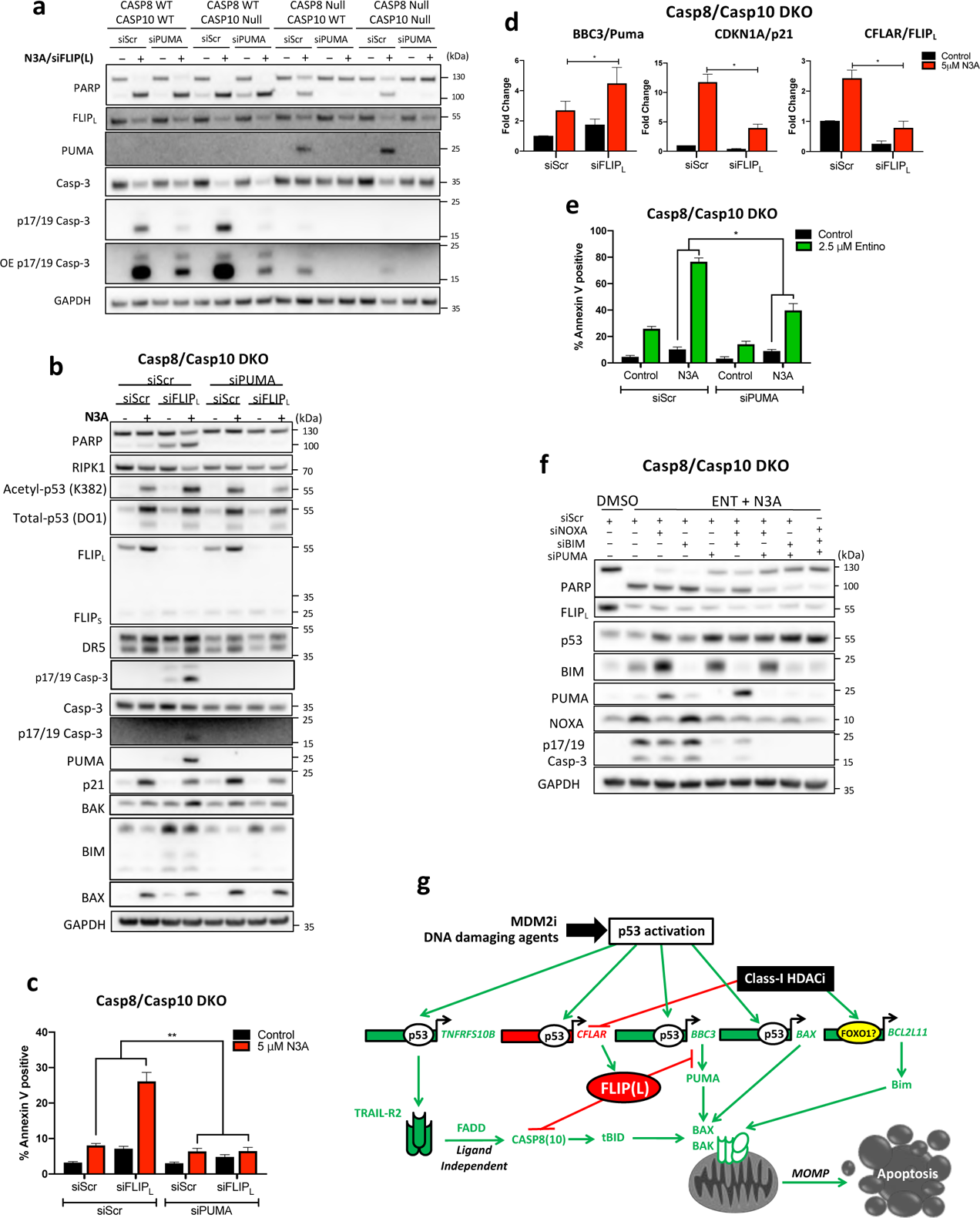
FLIP(L) inhibits p53-mediated upregulation of PUMA. (**a**) Western blot assessment of HCT116 Caspase-8/10 CRISPR single and dual KO (DKO) cells transfected for 24 h with 10 nM PUMA or control (Scr) siRNA prior to transfection with Scr/FLIPL siRNA for 6 h and treatment with 5 µM N3A for a further 48 h. Western blot (**b**) and Annexin-V/PI FACS (**c**) assessment of cell death in HCT116 caspase-8/10 DKO treated as in (**a**). (**d**) Annexin-V/PI FACS assessment of cell death in HCT116 caspase-8/10 DKO cells transfected with 10 nM PUMA or control (Scr) siRNA for 24 h prior to treatment with 5 µM Nutlin-3A (N3A) and 2.5 µM Entinostat (Entino) for a further 48 h. (**e**) Western blot assessment of HCT116 caspase-8/10 DKO cells transfected with 5 nM of the indicated siRNA’s for 24 h prior to treatment with 5 µM Nutlin-3A (N3A) and 2.5 µM Entinostat (Entino) for a further 48 h. (**f**) Quantitative RT-PCR of PUMA, p21 and FLIPL mRNA expression following transfection with Scr/FLIPL siRNA for 6 h and treatment with 5 µM N3A for a further 48 h. Data are normalized to RPL24 control for each sample and represented as mean ± s.e.m of four independent experiments. (**g**) Schematic representation of proposed mechanism of action. Error bars in (C/D/F) are represented as mean ± s.e.m. of at least three independent experiments. *P* values \**P* < 0.05; \*\**P* < 0.01; *** < 0.001 calculated by 2-way ANOVA (**c/d**) or Students t*-*test (**f**).

Silencing PUMA also significantly but incompletely rescued the cell death induced by Nutlin-3A/Entinostat in caspase-8/10-deficient cells (Fig. 7e/f). As has been previously reported (Inoue *et al*, 2007), another BH3-only protein BIM was significantly upregulated in response to Entinostat; moreover, we found that this induction was p53-independent (Supplementary information, Fig. S7h). Notably, co-silencing BIM alongside PUMA completely rescued the cell death induced by Nutlin-3A/Entinostat in caspase-8/10-deficient cells (Fig. 7e). The involvement of BIM in mediating the effects of Entinostat/Nutlin-3A likely explains the greater dependence of this combination on BAX than the siFLIP(L)/Nutlin-3A combination (Fig. 6i/j); BIM preferentially activates BAX, whereas BID preferentially activates BAK (Sarosiek *et al*, 2013). However, the resistance of p53-null cells to Entinostat/Nutlin-3A (Fig. 1/2) shows that this upregulation of BIM is insufficient to drive apoptosis in the absence of p53-mediated upregulation of other components of the apoptotic machinery. BIM upregulation was also observed in siFLIP(L)-treated caspase-8/10 deficient cells (Fig. 2b); however, this effect was abrogated in PUMA-depleted cells. Moreover, BIM upregulation in FLIP(L)-depleted cells was post-transcriptional (*data not shown*), suggesting that this effect may be a result of interplay between Bcl-2 family members, potentially caused by PUMA upregulation.

## Discussion

In the 50% of tumors harboring p53-WT, its function is typically suppressed; therefore, reactivation of p53’s latent tumor suppressive functions in these cancers is a highly attractive therapeutic approach (Brown *et al*, 2009). However, despite the development of selective p53 activating agents (best exemplified by MDM2 inhibitors, *e.g*. Nutlin-3A), none of these molecules have yet been clinically approved (Khoo *et al*, 2014). This potentially stems from the fact that these agents primarily induce cell cycle arrest rather than cell death in all but *MDM2*-amplified cancer cells (París *et al*, 2008; Tovar *et al*, 2006). Indeed, the mechanisms underlying the switch from p53-induced cell cycle arrest to p53-induced cell death remain surprisingly poorly understood, and strategies that enhance cell death induced by direct (*e.g.* MDM2 inhibitors) and indirect (*e.g.* DNA damaging agents) p53 activating agents are highly sought after.

HDACs have been shown to regulate acetylation of p53 in particular in its lysine-rich C-terminus, the hyper-acetylation of which has been correlated with enhancement apoptotic cell death (Knights *et al*, 2006; Mellert *et al*, 2011; Sykes *et al*, 2006; Brochier *et al*, 2013; Laptenko *et al*, 2015). HDAC2 has been reported to suppress p53 activity (Wagner *et al*, 2014) and is upregulated in CRC as a consequence of inactivation of the *adenomatous polyposis coli* (*APC*) tumour-suppressor gene, which is mutated in ∼80-85% of sporadic colorectal cancers (Zhu *et al*, 2004). Thus, we investigated the possibility that HDAC inhibition may enhance cell death induced by Nutlin-3A in CRC cells. Indeed, clear enhancement of CRC cell death was observed when Nutlin-3A was combined with the clinically relevant Class-I HDAC inhibitor Entinostat. Furthermore, we found that “non-acetylatable” p53 was unable to induce cell death in response to Nutlin-3A and Entinostat co-treatment. In contrast to our expectations, mRNA-Seq analyses revealed that Entinostat enhanced Nutlin-3A-induced expression of only one canonical p53 pro-apoptotic target, Noxa/*PMAIP1*. Paradoxically, we found that Entinostat *suppressed* Nutlin-3A-induced expression of a subset of apoptotic targets, most notably FLIP/*CFLAR*, which we identified as a novel direct p53 target gene and which our functional siRNA and CRISPR screens identified as a nodal suppressor of Nutlin-3A-induced cell death. Importantly, recent reports suggest that Class-I HDACi exert a significant proportion of their transcriptional modulatory effects through suppression of super-enhancer driven genes (Gryder *et al*, 2019), with maximal suppression requiring inhibition of HDAC1/2/3, consistent with our siRNA analysis, which showed that suppression of all 3 Class-I HDACs was required for maximal induction of Nutlin-3A-induced apoptosis (Fig. 1c).

We show that early acetylation-dependent p53 upregulation of FLIP(L) in response to Nutlin-3A prevents p53-mediated induction of apoptosis. This provides a novel mechanistic explanation for why p53 agonists fail to activate apoptosis in the majority of tumors despite robust upregulation of apoptosis target genes. While the caspase-8 inhibitory, anti-apoptotic effects of FLIP(S) are well-established, whether FLIP(L) is a promoter or inhibitor of caspase-8-mediated apoptosis is controversial (Riley *et al*, 2015). Herein, we clearly demonstrate that FLIP(L) upregulation in response to Nutlin-3A prevents caspase-8-mediated CRC cell death. Consistent with the canonical mechanism-of-action, the apoptosis induced in response to FLIP(L)-depletion in Nutlin-3A-treated cells is not only caspase-8-dependent, but also dependent on the adaptor protein FADD and the downstream effector BID, the cleavage of which promotes amplification of the cell death signal via the mitochondrial apoptotic pathway by activating BAX and (primarily) BAK. The upstream mechanism of action of caspase-8 was notable. Usually FADD-dependent recruitment of FLIP/caspase-8 to death receptors occurs in response to ligation of these receptors. However, we found that although TRAIL-R2 was clearly the primary upstream receptor primarily responsible for activation of caspase-8 in FLIP(L)-depleted Nutlin-3A-treated cells, its ligation by TRAIL was not required. Indeed, we detected a novel TRAIL-R2-FLIP(L) complex formed in response to Nutlin-3A, recruitment of caspase-8 into which was enhanced by FLIP(L)-depletion. Moreover, unlike the apoptosis induced by siFLIP(L)/Nutlin-3A, ligand (IZ-TRAIL)-induced apoptosis was only partially TRAIL-R2-dependent.

Notably, at later timepoints post Nutlin-3A treatment, significant levels of caspase-8-independent cell death were induced in FLIP(L)-depleted cells. We initially examined whether this was mediated by the caspase-8 and FLIP(L) paralog caspase-10. Similar to FLIP(L), the role of caspase-10 at the DISC is controversial, with it described as being able to compensate for lack of caspase-8 (Wang *et al*, 2001) unable to compensate for lack of caspase-8 (Sprick, 2002) and acting to inhibit caspase-8-mediated apoptosis (Horn *et al*, 2017b). Our results were interesting: in the presence of caspase-8, deletion of caspase-10 moderately enhanced cell death in siFLIP(L)/Nutlin-3A-treated cells (although this failed to reach significance); whereas in the absence of caspase-8, co-deletion of caspase-10 significantly decreased apoptosis induction. These results indicate that caspase-10 has a minor inhibitory role in caspase-8-proficient cells (potentially by interfering with procaspase-8 homo-dimerization, a prerequisite for caspase-8 activation, but can itself drive apoptosis in siFLIP(L)/Nutlin-3A-treated cells in the absence of caspase-8 (albeit less efficiently). We found that caspase-10-deficiency also enhanced IZ-TRAIL-induced apoptosis; however, unlike apoptosis induced by siFLIP(L)/Nutlin-3A, we found no evidence for caspase-10 being able to induce apoptosis in response to IZ-TRAIL in caspase-8-deificient HCT116 cells (Supplementary information, Fig. S4f). This difference between ligand-dependent and ligand-independent activation of TRAIL-R2 may be due to p53-mediated upregulation of caspase-10, which we found was induced in both RNA-seq and protein analyses along with FLIP in response to Nutlin-3A (Supplementary information, Table S1, HDAC visualiseR and Fig. 6b/d).

Even in the absence of caspase-8 *and* caspase-10, apoptosis was still observed in FLIP(L)-depleted, Nutlin-3A-treated cells. This cell death was also independent of TRAIL-R2, FADD and BID, but was BAX/BAK-dependent indicative of a novel role for FLIP(L) in regulating the intrinsic mitochondrial apoptotic pathway. Further analyses revealed an unexpected role for FLIP(L) in suppressing p53-mediated transcriptional upregulation of PUMA/*BBC3*, such that in siFLIP(L)/Nutlin-3A-treated cells, PUMA expression was upregulated and drove the apoptosis induced in caspase-8/10-deficient cells. Moreover, p53-induced p21 transcription was suppressed by siFLIP(L). Thus, in addition to its role in preventing ligand-independent TRAIL-R2-mediated apoptosis in response to p53 stabilization, these results are consistent with a novel pro-survival role for FLIP(L), *i.e.* suppression of p53-mediated upregulation of PUMA and promotion of p53-mediated upregulation of p21.

The mechanism of cell death induced by the combination of Nutlin-3A with Entinostat was almost identical to that induced by Nutlin-3A/siFLIP(L) with a couple of notable exceptions. As previously reported (Inoue *et al*, 2007) and confirmed by us, Entinostat significantly upregulates BIM; this upregulation has been suggested to be mediated by acetylation of FOXO1 (Yang *et al*, 2009). Along with PUMA, BIM was found to be responsible for the apoptosis induced by Nutlin-3A/Entinostat when the extrinsic apoptotic pathway was blocked by caspase-8/-10 co-deletion. As BIM preferentially activates BAX, and BID preferentially activates BAK (Sarosiek *et al*, 2013), this explains the differential dependence of the Nutlin-3A/Entinostat and Nutlin-3A/siFLIP(L) combinations on BAX and BAK respectively.

We speculate that acute concomitant induction of FLIP(L) and TRAIL-R2 in response to p53 stabilization enables epithelial cells to be *primed* to undergo apoptosis, whilst preventing induction of apoptosis *en masse*, which in epithelial cells would result in barrier dysfunction. Rapid commitment to cell death would then occur in response to secondary signaling event(s) (absent in cells treated with selective MDM2 inhibitors), which overcome FLIP(L)’s anti-apoptotic functions. In the context of more complex signals such as DNA damaging chemotherapy that result in oscillating levels of p53, its negative regulators and targets, modifications to p53 not induced by Nutlin-3A (such as acetylation) as well as activation/inactivation of additional signaling pathways may lead to down-regulation of FLIP(L) expression, thereby triggering commitment to apoptosis. These findings complement single cell studies that demonstrated the importance of late induction of inhibitors of apoptosis (IAP) family in raising the apoptotic threshold to block apoptosis in the context of sub-optimal p53 stabilisation by platinum (Paek *et al*, 2016). Furthermore, the specific and rapid upregulation of the long FLIP(L) splice form in response to p53 stabilisation may act not only to put the brakes on apoptosis, but also to inhibit the more pro-inflammatory RIPK1-dependent necroptotic cell death as the FLIP(L)-caspase-8 heterodimer can cleave RIPK1 (van Raam and Salvesen, 2012).

In summary, this work has uncovered novel, clinically-relevant biology, in which early p53-mediated upregulation of FLIP(L) prevents commitment of CRC cells to apoptosis by blocking activation of caspase-8/10 from a p53-induced ligand-independent TRAIL-R2 complex *and* by suppressing p53-mediated induction of PUMA (Fig. 7g). From a cancer therapeutics point of view, we show the potential of combining Nutlin-3A (and likely 2^nd^ generation MDM2 inhibitors) with Class-I HDAC inhibitors for the treatment of p53-WT CRC (and potentially other p53-WT cancers), and identify FLIP(L) as a critical p53-induced signaling node, the inhibition of which is necessary to promote apoptosis in response to MDM2 inhibition.

## Materials and Methods

### Reagents and Plasmids

The following reagents were used: SAHA (Vorinostat; Selleck Chemicals, Suffolk, UK), Nutlin-3a (MedChemExpress, Middlesex, NJ) and Entinostat (MS-275; Selleck Chemicals, Suffolk, UK). 5-Fluorouracil (5-FU) and Oxaliplatin were obtained as kind gifts from the Belfast City Hospital Cancer Centre. The WTp53, R248W and 8KR expression plasmids (pWZL-Blast vector) (Drosten *et al*, 2014) were kind gifts received from Mariano Barbacid, CNIO, Madrid.

## Antibodies

The following Western blot antibodies were used: PARP, Phospho Histone H2AX (S139), Phospho NF-κB p65/RelA (S536), HDAC2, Phospho IκBα (S32), IκBα, XIAP, BID, Caspase-3, Bax, Bim, RIPK1, TRAIL-R2/DR5 and α-tubulin (all Cell Signaling Technology); Acetyl p53 (K382), Acetyl p53 (K373), anti-p53 (mouse) (Abcam); anti-p53 DO-1 (total p53), PUMAα (PUMA), p21, GAPDH, HDAC1, HDAC3, NF-κB p65 (Total RelA), Bak, Noxa and FADD (all Santa Cruz Biotechnology); FLIP (AdipoGen); Acetyl Histone H3 (EMD Millipore); β-Actin (Actin) (Sigma-Aldrich); cIAP1, Caspase-8 (both Enzo Life Sciences); MDM2 (Oncogene Research Products); and Caspase-10 (MBL International). Human TRAIL/TNFSF10 antibody (R&D systems) was used at 100 ng/ml in neutralisation experiments.

### Western blot analyses

Western blot analyses were performed using samples lysed in RIPA Buffer supplemented with PhosSTOP (Roche) and Protease Inhibitor Cocktail (Roche). Equal amounts of protein (20-40 µg/well) were resolved by SDS-PAGE using the Novex system (Invitrogen) and transferred to nitrocellulose membranes using iBLOT (Invitrogen). Westerns were developed digitally using a FluorChem SP imaging system (Alpha Innotech) or the G:BOX ChemiXX6 gel doc system (Syngene).

### Immunoprecipitations

The DR5 DISC assay was performed as previously described (Majkut *et al*, 2014). For the post-lysis DR5 IP, cells were first lysed in 1ml DISC buffer (0.2% NP-40, 20 mM Tris-HCL (pH 7.4), 150 mM NaCl, 10% glycerol) for 90mins prior to the addition of 30 µl 4x AMG655 conjugated Dynabeads®. Samples were immunoprecipitated over night at 4^°^C. The AMG655-coated Dynabeads® were collected and washed 5 times in DISC buffer prior to resuspension in Laemmli buffer and analysis by Western blotting. Supernatants and inputs were also collected and analysed by Western blotting.

### Cell culture

HCT116 p53^+/+^, HCT116 p53^−/−^, HCT116 p53^−/R248W^cells and BAX^-/-^ (Sur *et al*, 2009) were kind gifts received from Prof Bert Vogelstein (Johns Hopkins University, Baltimore, MD). HCT116 *CASP8*^+/+^ and HCT116 *CASP8*^−/−^ cells (Paek *et al*, 2016) were kind gifts received from Prof Galit Lahav (Harvard Medical School, Boston, MA). HCT116 cells stably expressing the NFκB super-repressor and CT26 cells were a kind gift from Dr Aideen Ryan (NUI Galway, Ireland). HCT116 SMAC KO, DR5 KO and BAX/BAK DKO cells were a kind gift from Prof Markus Rehm (University of Stuttgart, Germany). HCT116 BID KO cells were a kind gift from Dr Lin Zhang, (University of Pittsburgh, Pittsburgh, PA).

LoVo and RKO cells were purchased from American Type Culture Collection (ATCC). HCT116 cell lines were cultured in McCoy’s 5A Medium supplemented with 10% fetal bovine serum, 50U/ml penicillin, 0.1mg/ml streptomycin, 2mM L-glutamine and 1mM sodium pyruvate (all Gibco). LoVo cell lines were cultured in Dulbecco’s Modified Eagle’s Medium (DMEM) (Gibco) supplemented with 10% fetal bovine serum, 50U/ml penicillin and 0.1mg/ml streptomycin. All cells were maintained at 37^0^C in a 5% CO2 humidified atmosphere and were regularly screened for the presence of mycoplasma using the MycoAlert Mycoplasma Detection Kit (Lonza, Basel, Switzerland).

### Stable cell lines

LoVo cells were transduced with retroviral pSUPER vectors expressing control or p53 shRNA under puromycin selection (0.5 µg/mL) as described previously (McDade *et al*, 2011). HCT116 p53-null cells were transduced with retroviral pWZL-Blast vectors expressing WT, R248W and 8KR p53 proteins and selected in 10 µg/mL Blasticidin (Sigma). HCT116 clones expressing a mutant form of the human IкB-α with serine 32 and 36 mutated into alanine (IкB-α SR) were generated using pRc-β-actin-IкB-α plasmid (SR) or a control pRc-β-actin plasmid (EV)(Luo *et al*, 2004) as previously described (Ryan *et al*, 2014).

### CRISPR Cell line generation

HCT116 FADD CRISPR cell lines were generated by retroviral infection with pLentiCRISPR with guide RNA targeting FADD; GCGGCGCGTCGACGACTTCG. Mixed population of cells were established following selection with 1 µg/ml puromycin. Cells were clonally selected for complete knockdown by Western blot. HCT116 caspase-8/caspase-10 double knockout cells were generated by retroviral infection of HCT116 caspase-8 CRISPR cells with pLentiCRISPRblast containing guide RNA targeting caspase-10:GACTGCTGCCCACCCGACAA. Mixed population of cells were established following selection with blasticidin (10 µg/ml). Cells were clonally selected for complete knockdown, verified by Western blot.

### RNA interference

Previously published siRNAs targeting TRAIL-R2/DR5, FLIP, FAS, FLIP(L) and FLIPS (Longley *et al*, 2005; Majkut *et al*, 2014); TRAIL-R1/DR4, (Wilson et al., 2009) RIPK1, Caspase-8 and a scrambled control (Crawford et al., 2018) were used. The siRNAs against HDAC1 and HDAC2 were commercially purchased (Qiagen). The siRNA against HDAC3 was described previously (Senese *et al*, 2007). The siRNA targeting p53 was purchased from Dharmacon (Dharmacon, Lafayette. CO) and has the following sequence: GUUCCGAGAGCUGAAUGAdTdT. ON-TARGETplus SMARTpool siRNA reagent targeting Caspase-10 (catalog # L-004402-00), PUMA (catalog # L-004380-00), NOXA (catalog # L-005275-00) and BIM (catalog # L-004383-00) were commercially purchased from Dharmacon. siRNA transfections were performed as previously described (Majkut et al., 2014).

### Apoptosis siRNA screen

siGENOME® SMARTpool® siRNA Library (Dharmacon) containing 4 sequences directed against each of 37 cell death related genes and controls were obtained in Echo-qualified 384-well polypropylene source plate (Labcyte, Sunnyvale, CA). siRNA pools were re-suspended to a final concentration of 2 µM and 150nL of 2 µM siRNA stocks were dispensed into 384-well assay plates using an Echo 525 Series Liquid Handler (Labcyte, Sunnyvale, CA), 10 µL of a 1:167 mix of RNAiMAX (Life Technologies) added, incubated for 1 h at room temperature, before addition of 900 cells per well in 20 µL of Optimem and incubated overnight (final concentration siRNA = 10 nM) The following day the transfection mixture was replaced with groWTh media with or without 2.5 µM Nutlin-3A. The plate was then incubated for 72 h at 37°C before measuring cell viability using CellTitreGlo reagent (Promega).

### Reverse transcription and quantitative RT-PCR analyses

Cells were homogenised using TriPure Isolation Reagent (Roche) and total RNA was then extracted using the PureLink RNA Mini Kit (Ambion). Alternatively, cells were lysed in buffer RLT plus (Qiagen) and RNA extracted using the RNeasy Plus extraction kit (Qiagen) as per manufacturer’s instructions. Subsequently, 2500ng of RNA was processed using the TURBO DNA-free Kit (Ambion) to remove contaminating genomic DNA, and 500ng of DNA-free RNA was used to generate cDNA using the High-Fidelity cDNA Synthesis Kit (Roche). Quantitative RT-PCR (RT-qPCR) was conducted using LightCycler 480 SYBR Green Master (Roche). The primers used are presented in Supplemental Supplementary Table4.

### Chromatin Immunoprecipitation (ChIP) and quantitative RT-PCR analyses

Cell fixation was carried out as previous described (McDade *et al*, 2014). Chromatin was isolated from 5 million cells and Chromatin immunoprecipitation for p53 was performed with DO-1 antibody (Santa Cruz Biotechnology) performed using the iDeal ChIP-seq kit for Transcription Factors according to manufacturer’s instructions (Diagenode) with direct method on the Diagenode IP-Star and purified with iPURE kit (Diagenode). Quantiitative PCR and fold enrichment estimated for CFLAR promoter estimated compared to negative control region (Supplementary Table 5) using as previously described (McDade *et al*, 2014).

### RNA-sequencing

Total RNA was extracted as above, and 1 µg of each RNA sample was used as a template to generate cDNA using the TruSeq Stranded mRNA Library Prep Kit (Illumina), following the manufacturer’s instructions for the low-sample (LS) protocol. These first-stranded cDNA samples were processed to barcoded cDNA libraries using an automated IP-Star (Diagenode), amplified, quantified and pooled and sequenced to yield 20-30M paired-end 75BP reads per sample on an Illumina Nextseq 500. FASTQ files were de-multiplexed, aligned to hg19 reference genome using BoWTie2 in using Basespace RNA-seq app (Illumina), resulting .bam files were mapped to RefSeq transcripts using Patek Genomics Suite (Partek) and gene-level RPKM normalized output used for differential gene expression analysis. Transcripts were filtered with an RPKM threshold of ≥0.5 in 50% of samples (12,411 genes) and significantly upregulated and downregulated transcripts identified using 3-way ANOVA with a mean RPKM fold-change threshold of ≥1.7x and False Discovery Rate (FDR) of 0.05. Resulting analyses can be visualised and downloaded from our Shiny app (functionalgenomicsgroup.shinyapps.io/HDAC_visualiseR/). Pathway analyses were conducted using the Enrichr online tool (Kuleshov *et al*, 2016). The data discussed in this publication have been deposited in NCBI’s Gene Expression Omnibus (Edgar *et al*, 2002) and are accessible through GEO Series accession number GSE113682. To review enter token uhwlougmbhglton into the box at: https://www.ncbi.nlm.nih.gov/geo/query/acc.cgi?acc=GSE113682

### CRISPR Screen

GeCKO Library V1 sgRNA library/pool targeting 17,419 genes (Shalem *et al*, 2014) was a kind gift from the Zhang laboratory. Library was amplified, lentivirus produced and titred as per Shalem et al. with following adaptations. Viral production: Six T175 flasks of HEK293T seeded at 40% confluence the day before of transfection. Media replaced with 13mL of Opti-MEM one hour prior to addition of transfection mix incubation for 6 hours then media replaced with fresh DMEM containing 10% FBS, 1% Pen/Strep supplemented with 10% BSA. X-tremeGENE HP (Roche), transfection agent was used at a ratio of 1:2 DNA:transfection reagent according to manufacturer’s instruction [per flask 15 µg of Gecko sgRNA pool, 7.7 µg of envelope plasmid pCMV-VSV-G (Addgene, cat # 8454), 11.5 µg of packaging plasmid psPAX2 (Addgene, cat # 10668)]. Supernatant was collected after 60 hours and virus purified by Lent-X Maxi Purification kit (Clontech), eluted and resuspended in DMEM supplemented with 10% BSA, aliquoted and stored at −80°C. Twenty million HCT116 p53 +/+ and -/- cells were transduced at MOI of 0.3 with Gecko v1 Lentivirus for 72 hours and selected for 8 days in puromycin. 30×10^6^ cells were harvested (representing an estimated coverage of 300x cells) and 30×10^6^ cells plated for treatment the next day with 5 µM Nutlin-3A, DMSO vehicle, or media only for a further 11 days. Cells were passaged when reached 70-80% confluence again maintaining 30×10^6^ cells at each serial passage, Nutlin-3A added in fresh media every 3 days. DNA was extracted from cells with Blood & Cell Culture DNA Maxi Kit of Qiagen according to manufacturer’s instruction with the final DNA extracted calculated to have a coverage ratio between 110X to 300X. PCRs sgRNA inserts were amplified as per Shalem et al. using Herculase II Fusion DNA Polymerase (Agilent, cat # 600675). Two step amplification was carried out with 18 cycles with primers to amplify LentiCRISPR sgRNA (Tm at 55°C) and 24 cycles for the second with Illumina adaptor and barcoded staggered adapters (Tm at 60°C). Resulting ∼350BP PCR amplicons were size selected using SPRI select (Beckman Coulter), quantified, pooled and sequenced on Nextseq 500 single end 80Bp read plus 8Bp index. Adapters were trimmed from de-multiplexed FASTQ files using Cutadapt (https://pypi.python.org/pypi/cutadapt) to extract sgRNA sequences which were aligned to the Gecko V1 design file and count tables generated with MAGeCK software V5. Positively and negatively selected sgRNAs were identified at sgRNA, gene and pathway level using MAGeCK RRA algorithm (Li *et al*, 2014)

### Flow cytometry analyses

Cells were harvested, and Annexin V and propidium iodide (PI) positivity analysed as previously described (Majkut *et al*, 2014). Analysis was carried out on either the BD LSR-II flow cytometer (BD Biosciences) or the BD Accuri^TM^ C6 flow cytometer (BD Biosciences).

### High-content microscopy

Depending on time point and cell type 2-8,000 cells per well were seeded onto black 96-well glass-bottom plates (*In Vitro* Scientific) in a volume of 100 µL/well prior to treatment. After treatment, 50 µL of cell staining solution (3 µg/mL FITC-tagged Annexin V (BD Biosciences), 1 µg/mL Propidium Iodide (PI) (Sigma Aldrich) plus 30 µg/mL Hoechst 33342 (Invitrogen) in PBS) was added to each well. Plates were then incubated in the dark for 15 minutes at room temperature prior to reading on an ArrayScan XTI Live High Content Platform (ThermoFisher). Hoechst nuclear stain was used to locate single-cells prior to measuring individual fluorescence intensities for PI, Annexin V or dual positivity. Control wells containing positive/negative populations for apoptosis were used to set a threshold for scoring individual cells as ‘live’ or ‘dead’ based on average PI or Annexin V fluorescence intensity. This binary score was applied to the entire plate to generate a readout of % Annexin V, PI or AnnexinV/PI positive cells per well from a total of 2,000 assessed cells.

### Cell viability assay

Cell viability was assessed using the Cell Titre-Glo® assay according to manufacturer’s instructions (Promega, Madison, WI, USA).

### In vivo studies

All procedures were carried out in accordance with the Animals (Scientific Procedures) Act, 1986 in compliance with institutional guidelines and authority regulations under SPF conditions. For transplant studies, 8-12 week old female BALB/c mice received a single, subcutaneous injection of 1×10^6^ CT26 cells (ATCC) in PBS (Invitrogen). Tumour volumes (mm^3^) were measured twice weekly using the formula: *1/2(a x b^2^)*, where *b* is the shorter measurement. Once tumours were established (average volume 100mm^3^), animals were randomised and assigned to four treatment groups: Vehicle (30% cyclodextrin (Entinostat vehicle, OG)/PBS (FOLFOX vehicle, IP), FOLFOX alone (10mg/kg 5-FU + 1mg/kg Oxaliplatin (IP), 30% cyclodextrin (OG)), Entinostat alone (10mg/kg Entinostat (OG), PBS (IP)) or a combination of both (10mg/kg 5-FU + 1mg/kg Oxaliplatin (IP), 10mg/kg Entinostat (OG). For FOLFOX treatment, 5-FU was administered daily and Oxaliplatin on Day 1 of 7 only. Mice were treated for 3 or 7 days, with tumour volume assessed throughout, and tumours harvested and fixed for FFPE processing at endpoint.

### TUNEL Assay

Terminal deoxynucleotidyl transferase dUTP nick end labelling (TUNEL) assay (ApopTag Fluorescein Direct InSitu Apoptosis Detection Kit, Millipore) was used to quantify apoptosis in CT26 allografts following 72 hour treatment. Briefly, tumours were excised and fixed for 16hrs in 10% Neutral-Buffered formalin (Atom Scientific) before paraffin-embedding and sectioning. 5 µm sections were processed for TUNEL staining according to manufacturer’s instructions and coverslips mounted using Vectashield Antifade Mounting Medium with DAPI (H-1200), and allowed to set in the dark at room temperature overnight. The following day, DAPI/FITC images from 3 fields of view (FOV) per tumour were captured using the Zeiss Axiovert 200M at 20x magnification, and the number of FITC (TUNEL) positive cells per FOV was counted. Staining and counting was repeated to validate results.

## Statistical Analysis

Statistical analyses were performed using Prism 8.0 software (Graphpad). Experimental results were compared using a two-tailed Students t-test, One-Way ANOVA with multiple comparisons and Bonferroni Post-test, Two-Way ANOVA or Three-Way ANOVA where appropriate; *p ≤ 0.05 **p ≤ 0.01 ***p ≤ 0.001 and ****p ≤ 0.0001. Experiments were replicated at least 2 times and in most cases 3 or more.

## Acknowledgments

Work in this manuscript was primarily supported by studentships (PI: SSM) sponsored by Northern Ireland DfE and DEL. As well as funding from CRUK Program grants C11884/A24367 (PI: DBL; CI: SSM) and C212/A13721(PI: PGJ, Co-I DBL), ECMC C36697/A25176, RCUK | Biotechnology and Biological Sciences Research Council (BBSRC) - BB/T002824/1 (PI McDade, Longley). RNA-seq data generation in this study was supported by the “Queen’s FMHLS Genomics Core Technology Unit”

## Author Contributions

Conceptualization, SSM, DBL, PGJ.; Methodology, SSM, DBL, AM, AL, NC, AER, LJE; Investigation, AM, AL, NC, FF, TS, GG, KM, CH, PG; Writing – Original Draft, SSM, DBL; Writing – Review & Editing, PD; Funding Acquisition, SSM, DBL, PGJ; Resources, GQ, MW, WLA; Visualization SSM, DBL, AM, AL, MW. We thank Professor Bert Vogelstein (Johns Hopkins University School of Medicine, Baltimore, MD) for kindly gifting the HCT116 p53^+/+^ and p53^-/-^ cell lines, Professor Markus Rehm (University of Stuttgart, Germany) for supplying the HCT116 DR5 KO, BAX/BAK DKO and SMAC KO cells and Professor Galit Lahav (Department of Systems Biology, Harvard Medical School, Boston, MA) for providing the HCT116 caspase-8 CRISPR cells. We also thank Dr Lin Zhang (University of Pittsburgh, Pittsburgh, PA) for supplying the HCT116 BID KO cells.

---

**Supplementary Fig. 1.**
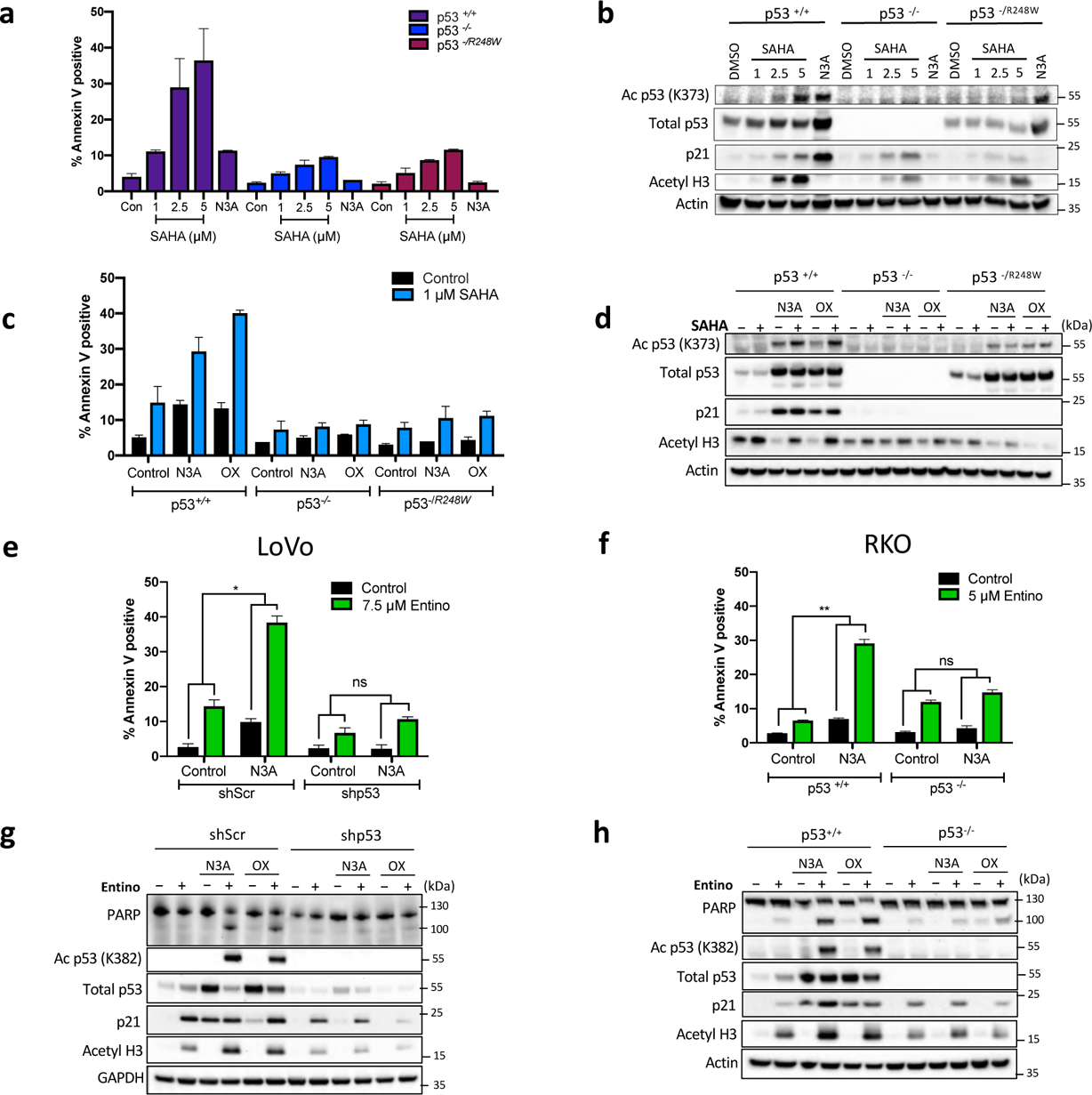
HDAC inhibition enhances p53-dependent apoptotic cell death. Assessment of cell death with Annexin-V/PI FACS (a) and Western blot analysis (b) in p53-WT, null and R248W mutant HCT116 isogenic cell lines treated with 1, 2.5 or 5 µM SAHA or 5 µM Nutlin-3A (N3A) for 24 h. Annexin-V/PI FACS (c) and Western blot (d) analyses of p53-WT, null and R248W mutant HCT116 isogenic cell lines treated with 5 µM N3A or 1 µM Oxaliplatin (Ox) for 24 h prior to treatment with 1 µM SAHA for a further 24 h. (e) Annexin-V/PI FACS cell death analysis of LoVo cell lines stably transduced with shRNA targeting p53 (shp53) or scrambled control (shScr) treated with 5 µM N3A for 24 h prior to treatment with 7.5 µM Entino for a further 72 h. (f) Annexin-V/PI FACS in RKO p53-WT and null isogenic cell lines treated with 2.5 µM N3A for 24 h prior to treatment with 5 µM Entino for a further 24 h. (g/h) Western blot analysis of LoVo and RKO isogenic cell lines treated as in (e/f). Error bars in (d/e) are represented as mean ± s.e.m. of at least three independent experiments. *P* values \**P* < 0.05; \*\**P* < 0.01, \*\*\**P* < 0.001 calculated by 2-way ANOVA.

**Supplementary Fig. 2.**
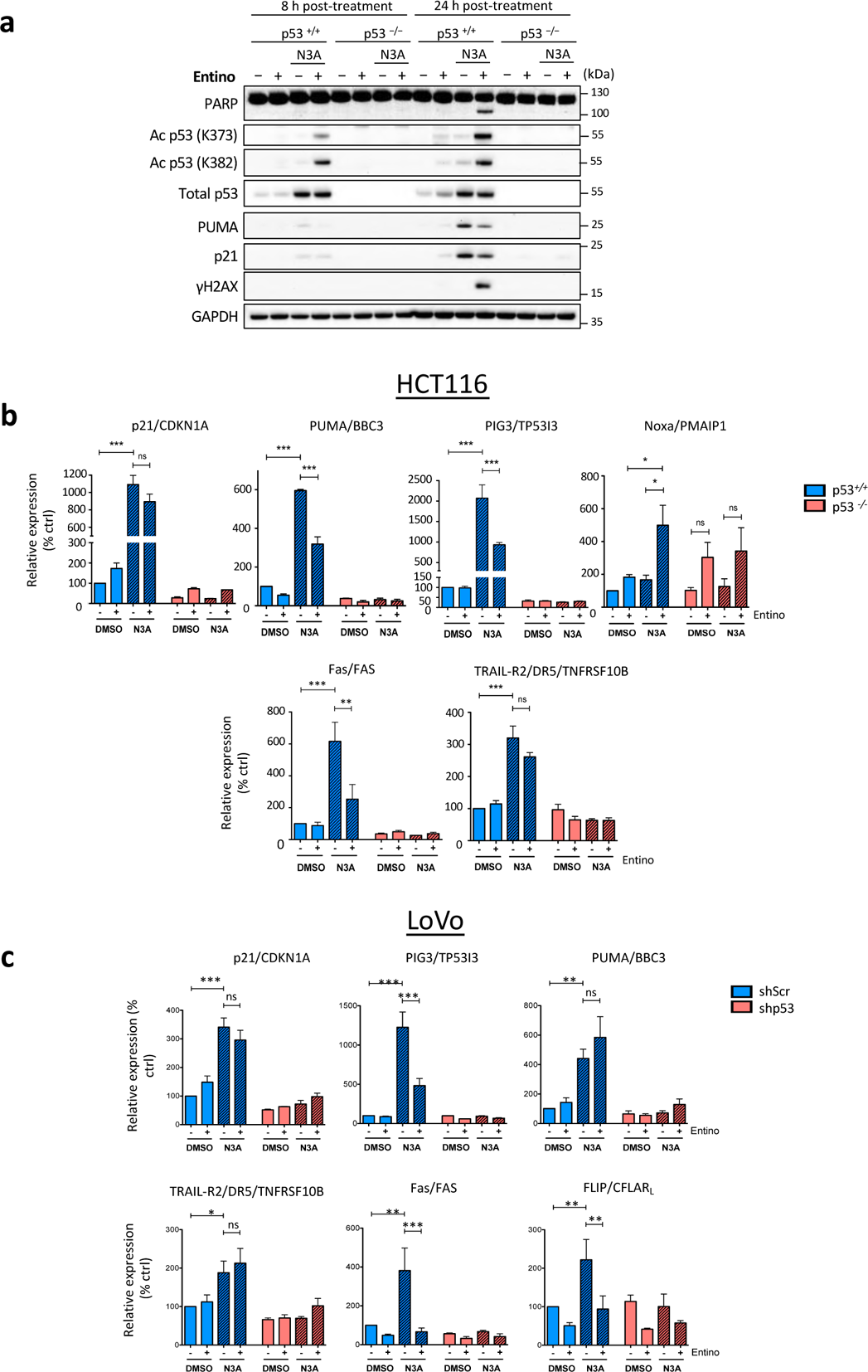
Analysis of impact of Entinostat Nutlin-3A in HCT116 p53-WT and Null models. (**a**) Western blot analysis of p53-WT and null HCT116 cells treated with 5 µM Nutlin-3A (N3A), 2.5 µM Entinostat (Entino) or the combination of both agents for 8 or 24 h. (**b**) Quantitative real-time PCR analysis of p53 target gene expression p53-WT and null HCT116 cells treated with 5 µM Nutlin-3A (N3A), 2.5 µM Entinostat (Entino) or their combination for 8 h. Expression values are normalized to RPL24 control for each sample. (**c**) Quantitative real-time PCR analysis of p53 target gene expression in LoVo cells stably expressing scrambled or p53 shRNA following treatment with 2.5 µM Nutlin-3A, 7.5 µM Entinostat or their combination for 24 h. Expression values are normalized to RPL24 control for each sample. Error bars in (**b/c**) are represented as mean ± s.e.m of at least three independent experiments. *P* values \**P* < 0.05; \*\**P* < 0.01, \*\*\**P* < 0.001 calculated by paired samples 1-way ANOVA with Bonferroni post-tests.

**Supplementary Fig. 3.**
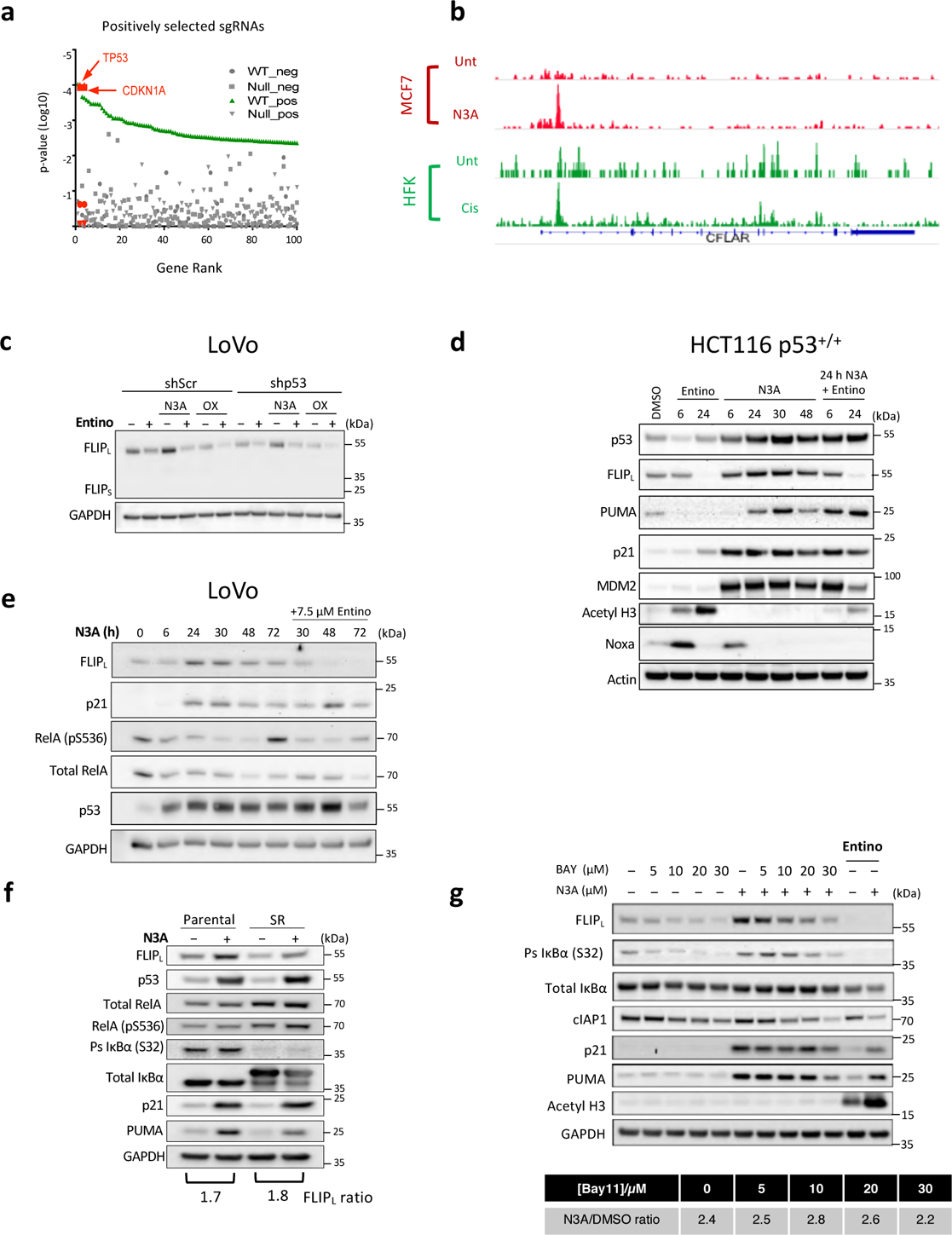
Functional genomics analyses identify *CFLAR*/FLIP as a direct HDAC1/2/3-dependent pro-survival target of p53. (**a**) Results of a genome-wide CRISPR screen for enhancers of Nutlin-3A sensitivity or resistance. The *p*-value for MAGeCK gene level analysis of top 100 genes whose CRISPR guide RNAs (positively (resistance) selected CRISPR guide RNAs (sgRNAs) are depicted, highlighting p53 (*TP53*) and p21 (*CDKN1A*). (**b**) ChIP-seq tracks for p53 binding to the FLIP/CFLAR promoter I MCF7 cells treated with Nutin-3A or HFKs treated with Cisplatin. (**c**) Western blot analysis of LoVo cell lines stably expressing scrambled or p53 shRNA treated with 2.5 µM N3A or 1 µM Oxaliplatin (OX) for 24 h prior to treatment with 7.5 µM Entino for a further 72 h. (**d**) Western blot analysis of p53-WT HCT116 cells treated with 5 µM N3A with or without 2.5 µM Entino for the indicated times. (**e**) Western blot analysis of LoVo cells treated with 2.5 µM N3A alone or with 7.5 µM Entino for the indicated times. Western blot and densitometric analysis of the ratio of FLIP induction in (**f**) HCT116 IkB-super-repressor (SR) and matched parental cells treated with 5 µM N3A for 24 h and (**g**) p53-WT HCT116 cells treated with 5 µM N3A in the presence and absence of the NFkB inhibitor BAY117082 or 2.5 µM Entino for 24 h.

**Supplementary Fig. 4.**
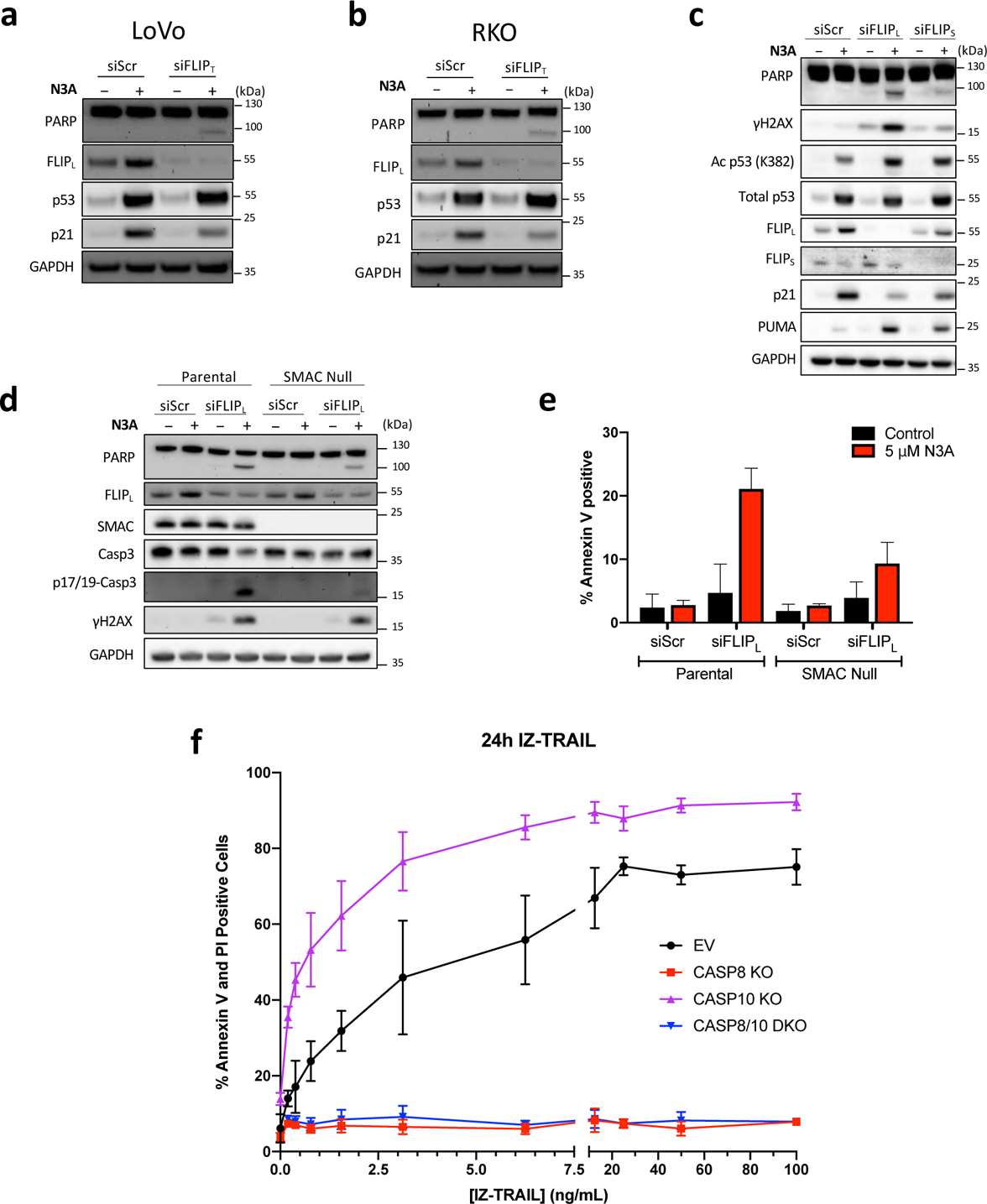
Nutlin-3a induces a dependence on FLIP. Western blot assessment of cell death in LoVo (**a**) and RKO (**b**) p53-WT cells transfected for 6 h with 10 nM scrambled (siScr) or a FLIPT specific siRNA prior to treatment with 5 µM Nutlin-3A (N3A) for a further 24 h. (**c**) Western blot analysis of HCT116 p53-WT cells transfected with 10 nM FLIPL- or FLIPS-specific siRNAs or control siRNA for 6 h prior to treatment with 5 µM N3A for a further 24 h. (**d**) Western blot and (**e**) Annexin-V/PI FACS assessment of cell death in HCT116 SMAC null and matched parental cells following treatment with 10 nM FLIPL or control siRNA for 6 h prior to treatment with 5 µM N3A for a further 24 h. (**f**) High Content Microscopy analysis of cell death in HCT116 control (EV) and Caspase-8- and Caspase-10-knockout (KO) and Caspase-8/10-double knockout (DKO) daughter cells 24 h after treatment with IZ-TRAIL. Error bars in (**b**) are represented as mean ± s.e.m. of at least three independent experiments. *P* values for combined treatments \**P* < 0.05; \*\**P* < 0.01, \*\*\**P* < 0.001 calculated by 2-way ANOVA. Error bars in (**e/f**) are represented as mean ± s.d. of two independent experiments.

**Supplementary Fig. 5.**
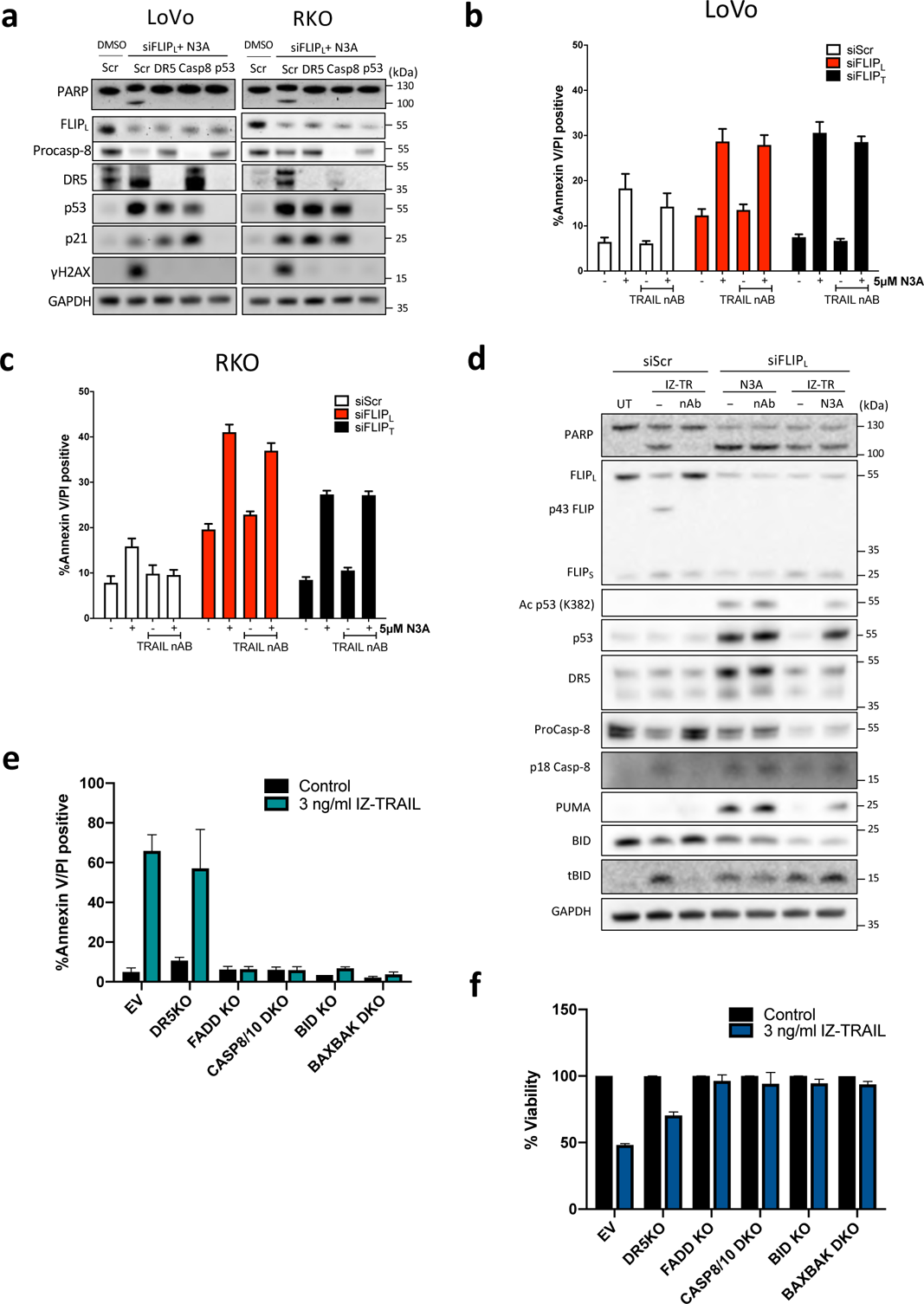
Nutlin-3A primes p53 wild-type CRC cells for TRAIL-independent cell death. (**a**) Western blot assessment of LoVo and RKO cells transfected with the indicated siRNA’s (10 nM) for 24 h followed by transfection with siFLIPL for 6 h and treatment with 5 µM Nutlin-3A (N3A) for a further 24 h. (**b**) High Content Microscopy analysis of cell death in LoVo cells transfected with the indicated siRNA’s for 6 h prior to treatment with 5 µM N3A +/-100 ng/ml TRAIL-neutralising antibody (TRAIL nAB) for 48 h. (**c**) High Content Microscopy analysis of cell death in RKO cells treated as in (**b**). (**d**) Western blot analysis of the effect of TRAIL neutralizing antibody treatment (100 ng/ml) on cell death in HCT116 cells transfected for 6 h with 10 nM scrambled (siScr) or FLIPL siRNA prior to treatment with 3 ng/ml isoleucine zipper TRAIL (IZ-TR) and/or 5 µM Nutlin-3A (N3A) for a further 24 h. (**e**) High Content Microscopy analysis of cell death in HCT116 control (EV) and TRAIL-R2/DR5-, FADD- and BID knockout (KO) and Caspase-8/10- and BAX/BAK double knockout daughter cells 24 h after treatment with 3 ng/ml IZ-TRAIL. (**f**) CellTiterGlo cell viability analysis of HCT116 control (EV) and TRAIL-R2/DR5-, FADD- and BID KO and Caspase-8/10- and BAX/BAK DKO daughter cells 24 h after treatment with 3 ng/ml IZ-TRAIL. Error bars in (**b/c/e/f**) are represented as mean ± s.d. of two independent experiments.

**Supplementary Fig. 6.**
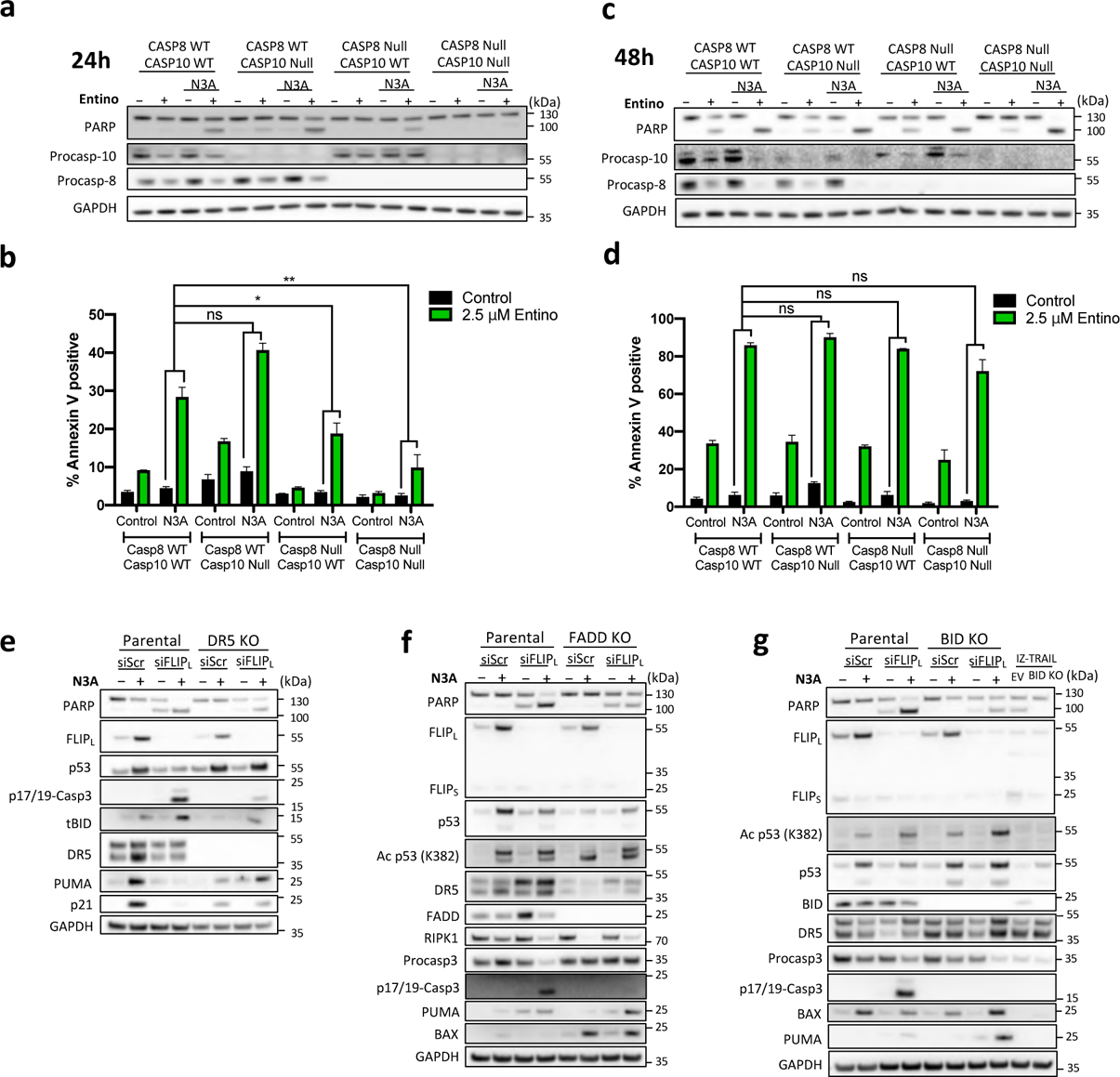
Death proceeds independently of extrinsic pathway components at later timepoints. Western blot (**a/c**) and Annexin-V/PI FACS analysis of cell death (**b/d**) in HCT116 Caspase-8/10 CRISPR single and dual KO cells treated with 2.5 µM Entinostat (Entino) and 5 µM Nutlin-3a (N3A) for 24 h (**a/b**) and 48 h (**c/d**). Western blot assessment of cell death in HCT116 isogenic for (**e**) DR5/TRAIL-R2, (**f**) FADD and (**g**) BID knockout transfected for 6 h with 10 nM scrambled (siScr) or FLIPL siRNA prior to treatment with 5 µM Nutlin-3A (N3A) for a further 48 h. 3 ng/ml IZ-TRAIL acts as a positive control for BID KO. Error bars in are represented as mean ± s.e.m. of at least three independent experiments. *P* values \**P* < 0.05; \*\**P* < 0.01, \*\*\**P* < 0.001 calculated by 2-way ANOVA.

**Supplementary Fig. 7.**
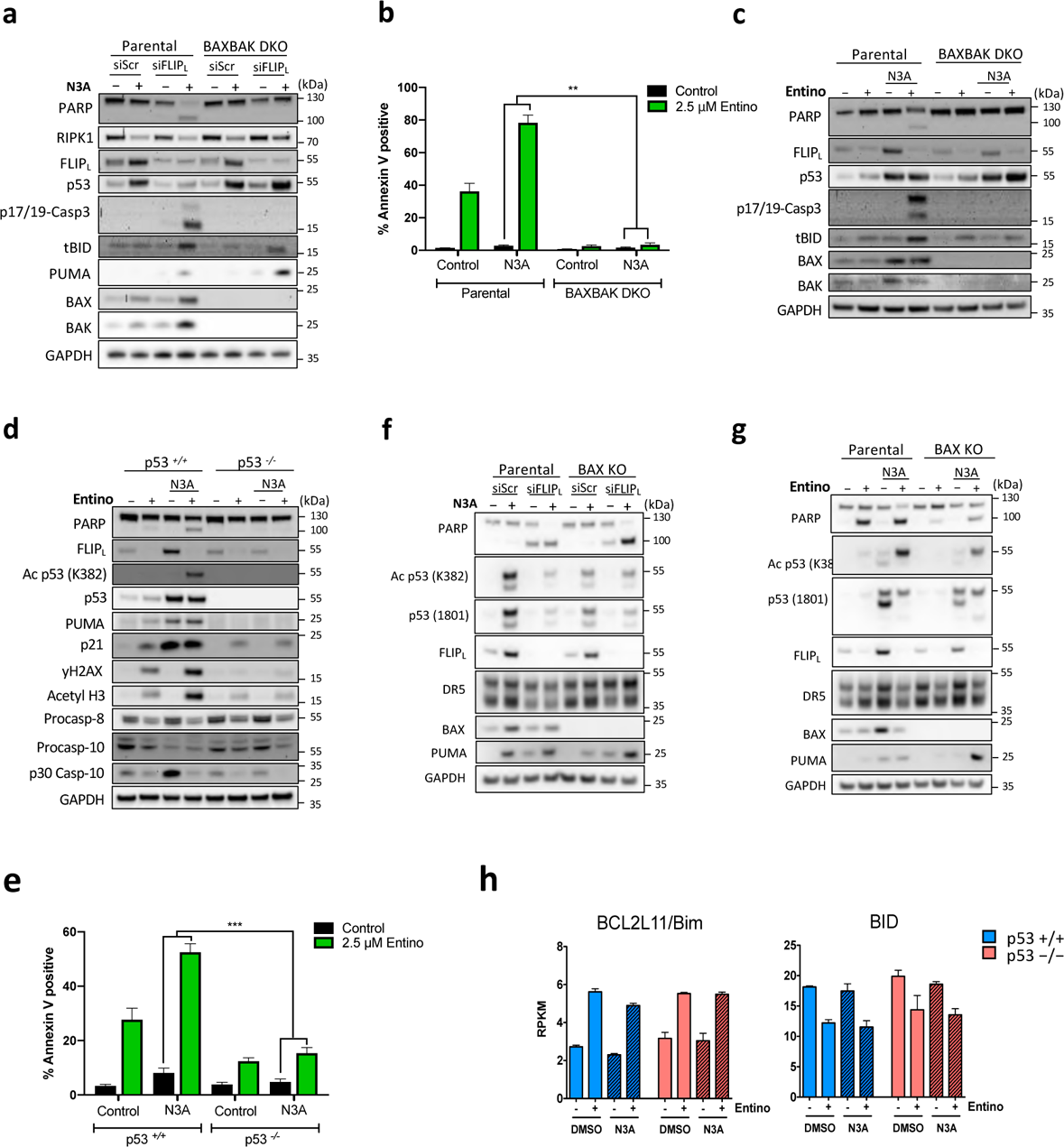
Late death is driven by the mitochondria. (**a**) Western blot assessment of cell death in HCT116 BAXBAK double knockout (DKO) and matched parental cells transfected for 6 h with 10 nM scrambled (siScr) or FLIPL siRNA prior to treatment with 5 µM Nutlin-3A (N3A) for a further 48 h. (**b**) Annexin-V/PI FACS and (**c**) Western blot analysis of cell death in BAXBAK DKO and matched parental cells treated with 2.5 µM Entinostat (Entino) and 5 µM Nutlin-3a (N3A) for 48 h. Western blot (**d**) and Annexin-V/PI FACS (**e**) analysis of cell death in HCT116 p53-WT and null cells treated with 2.5 µM Entinostat (Entino) and 5 µM Nutlin-3a (N3A) for 48 h. Western blot analysis of BAX KO and matched parental cells treated with either (**f**) 10 nM scrambled (siScr) or FLIPL siRNA for 6 h prior to treatment with 5 µM Nutlin-3A (N3A) for a further 48 h or (**g**) 2.5 µM Entinostat (Entino) and 5 µM Nutlin-3a (N3A) for 48 h. (**h**) RPKM values for *BCL2L11*/BIM and BID from mRNA-seq performed in HCT116 p53-WT and null isogenic cells (described in Fig. 2(**c**)) represented as mean RPKM ± s.d. of two independent experiments. Error bars in (**b/e**) are represented as mean ± s.e.m. of at least three independent experiments. *P* values \**P* < 0.05; \*\**P* < 0.01, \*\*\**P* < 0.001 calculated by 2-way ANOVA.

